# Specific Pupylation as IDEntity Reporter (SPIDER) for the identification of Protein-Biomolecule interactions

**DOI:** 10.1101/2022.05.26.493537

**Authors:** He-Wei Jiang, Hong Chen, Yun-Xiao Zheng, Xue-Ning Wang, Qingfeng Meng, Jin Xie, Jiong Zhang, ChangSheng Zhang, Zhao-Wei Xu, ZiQing Chen, Lei Wang, Wei-Sha Kong, Kuan Zhou, Ming-Liang Ma, Hai-Nan Zhang, Shu-Juan Guo, Jun-Biao Xue, Jing-Li Hou, Zhe-Yi Liu, Wen-Xue Niu, Fang-Jun Wang, Tao Wang, Wei Li, Rui-Na Wang, Yong-Jun Dang, Daniel M. Czajkowsky, Yu Qiao, Jia-Jia Dong, JianFeng Pei, Sheng-Ce Tao

## Abstract

Protein-biomolecule interactions play pivotal roles in almost all biological processes, the identification of the interacting protein is essential. By combining a substrate-based proximity labelling activity from the pupylation pathway of *Mycobacterium tuberculosis*, and the streptavidin (SA)-biotin system, we developed <u>S</u>pecific <u>P</u>upylation as <u>IDE</u>ntity <u>R</u>eporter (SPIDER) for identifying protein-biomolecular interactions. As a proof of principle, SPIDER was successfully applied for global identification of interacting proteins, including substrates for enzyme (CobB), the readers of m^6^A, the protein interactome of mRNA, and the target proteins of drug (lenalidomide). In addition, by SPIDER, we identified SARS-CoV-2 Omicron variant specific receptors on cell membrane and performed in-depth analysis for one candidate, Protein-g. These potential receptors could explain the differences between the Omicron variant and the Prototype strain, and further serve as target for combating the Omicron variant. Overall, we provide a robust technology which is applicable for a wide-range of protein-biomolecular interaction studies.

## Introduction

It is known that the vast majority of biomolecules carry out their biological functions through interacting with proteins. There is thus tremendous interest to identify the global set of protein-biomolecule interactions, like protein-protein interactions (PPIs), protein-nucleic acids interactions (PNIs) and protein-small molecule interactions (PSMIs).

PPIs are fundamental for most biological processes, such as the formation of cellular structures and enzymatic complexes or signaling pathways. Traditional methods for PPI identification include immuno-precipitation (IP)^1^, GST-pull down^2^, protein microarray^3^, and affinity purification mass spectrometry (AP-MS)^4^. For these methods, the binding between the bait protein and the target protein is usually not covalent, and thus these methods are not well suited for the identification of weak and transient binders or applications where stringent washing is required, as is the case when studying ligand-cell surface receptor interactions. Proximity labeling methods, such as BioID^5^, APEX^6^, and PUP-IT^7^, are attractive alternative choices for the identification of PPIs^8^. However, the protein of interest is usually required to be fused to an enzyme with proximity-tagging activity. While this does afford the possibility of labeling inside cells, the efficacies of these proximity tagging methods are highly dependent on the proper expression of the enzyme. In addition, the size of the proximity tagging enzyme is typically not small (27 to 54 kDa)^5–7^, which limits the proteins that could be studied.

DNA binding proteins (DBPs) are a ubiquitous family of proteins involved in many fundamental biological processes^9^. RNA binding proteins (RBPs) are also widely abundant and known to play a number of essential roles in cells^10,11^. Traditional nucleic acid-centric methods for the identification and characterization of DBPs or RBPs include oligo-capture pulldown and protein microarrays^12,13^. For these methods though, sophisticated operations include UV cross-linking are needed to balance efficiency and specificity. Alternately, a number of proximity labeling methods have been developed, such as CAPTURE and CASPEX for DNA^14,15^ and CRUIS and CARPID for RNA^16,17^. For these methods, the nucleic acid of interest is usually targeted by a dCas protein, fused with a proximity-based tagging enzyme, and guided by sgRNA. These methods require highly specific sgRNA that recognizes the target sequence, sgRNA off-target is a concern^18^. Another limitation is the large size of dCas protein, which may sterically interfere with PNI^8^. In addition, the existing proximity labeling methods are difficult to identify specific modified nucleic acids-protein interaction because it is hard to modified nucleic acids at special sites *in vivo*.

Small molecules, defined as low molecular weight organic compounds (typically <1,000 Da), which could be either natural or artificial, are widely applied as tools in scientific research, and extensively used for therapeutics^19–22^. Many of these small molecules are believed to carry out their function as a consequence of interactions with specific proteins (that is, protein-small molecule interactions (PSMIs)). Thus, for any small molecule of interest, identification of its target protein(s) is the key to understand its mode-of-action (MOA). Affinity purification mass spectrometry (AP-MS) is widely used for small molecule target identification^23,24^. However, usually, the binding between the bait small molecule and the target protein is not covalent, and thus, it is challenging to balance the sensitivity and specificity, as a gentle wash might not effectively remove all non-specific binding proteins, while a stringent wash may result in loss of genuine targets. To address this issue, new approaches have been explored. For example, covalent affinity probes (also known as tri-functional affinity probes) have been shown to effectively identify low abundant targets through additional covalent capturing^25,26^. However, the small molecules are required to be equipped with additional photo-reactive or electrophilic groups in a case-by-case manner through tedious *de novo* synthesis^27,28^.

Overall, there are varied limitations of the current methods for the identification of protein-biomolecule interactions. This situation severely hampers our understanding of the underlying molecular mechanisms. Herein, we serendipitously discovered a substrate-based proximity labelling activity from the pupylation pathway of *Mycobacterium tuberculosis*. Taking advantage of this activity, we developed <u>S</u>pecific <u>P</u>upylation as <u>IDE</u>ntity <u>R</u>eporter (SPIDER) as an efficient and widely applicable method for the identification of protein-biomolecule interactions. We demonstrated the ease and wide-ranging ability of SPIDER by characterizing a variety of challenging protein-biomolecule interactions, from PPI, PNI, PSMI to ligand-receptor interactions on cell surface.

## Results

### Design and validation of the SPIDER proximity-tagging system

In *Mycobacterium tuberculosis*, PafA catalyzes the covalent linkage of the C-terminus of Pup^E^ to target proteins for proteasomal degradation^29^. We serendipitously discovered that PafA can covalently link Pup^E^ to its proximal proteins when Pup^E^ is attached to a protein of interest via its N-terminus (Extended Data Fig. 1a to g). This proximal ligation involves the attachment of the substrate (Pup^E^) to protein of interest, which is distinct from the known proximity ligation reaction using PafA fused protein^7^. Based on this discovery, we designed SPIDER to identify protein-protein interactions (Fig. 1a). A key component of SPIDER is Pup^E^ fused streptavidin (SA-Pup^E^) (Fig. 1a and Extended Data Fig. 1h). The distinguishing feature of SA-Pup^E^ is that pup maintains a disordered structure in the absence of cofactor binding^30^, which provides a large degree of flexibility of potential linking configurations to the prey protein. In the SPIDER assay, the bait is firstly biotinylated using any of the well-established methods^31^. The proteins (the prey) that bind to the bait are then covalently linked to the Pup^E^ peptides in the SA-Pup^E^ tetramer by PafA (Fig. 1a and Extended Data Fig. 1h). For assays involving unknown prey proteins, the binding reaction can be simply visualized using gel electrophoresis to monitor a shift in the mass of the SA-Pup^E^. Alternatively, biotin agarose can be used to enrich the SA-Pup^E^ linked proteins, followed by MS identification. The biotin binding activity of SA-Pup^E^ retains the stability as wild type even in 8 M urea (Extended Data Fig. 1i).

**Fig.1.**
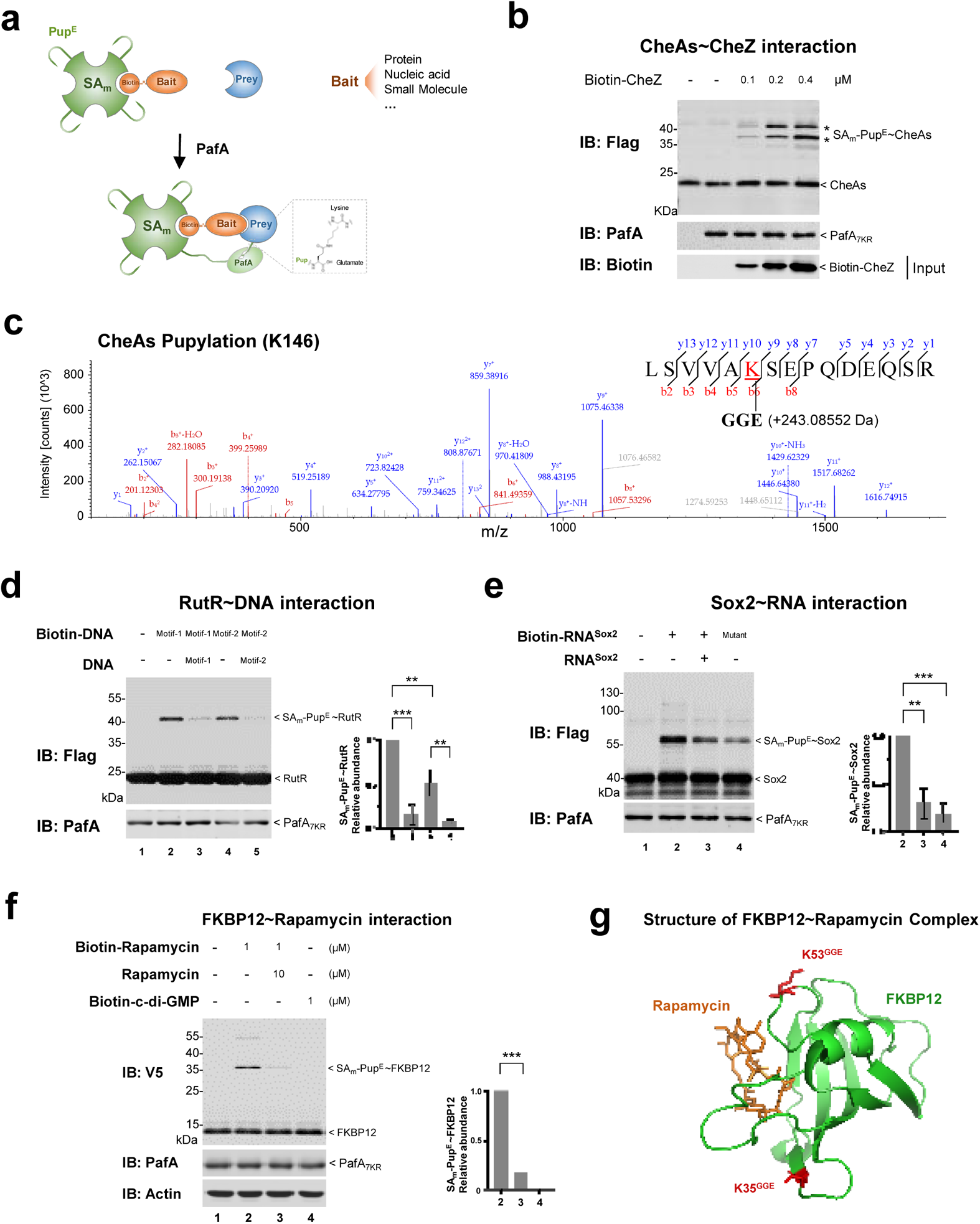
Design and validation of the SPIDER proximity-tagging system. **a,** Schematic design of SPIDER. The key components of SPIDER are Pup^E^ fused streptavidin (SA-Pup^E^) and PafA. When a biotinylated bait including protein, nucleic acid and small molecules binds to a prey protein, it brings pup^E^ to the proximity of the bait through the binding of biotin and SA-pup^E^. PafA is recruited and covalently links pup^E^ to the adjacent lysine/s on the prey protein, thus transform the binding of the biotinylated bait and the prey protein into a covalent linkage between SA-pup^E^ and the prey protein. The SA-Pup^E^-prey protein complex is enriched by biotin agarose through stringent wash, *e. g*., 8 M urea. The binding of the biotinylated bait and the prey protein is visualized through gel electrophoresis or identified by mass spectrometry. **b,** SPIDER assays with Flag-tagged CheAs and biotin-CheZ (WT). **c,** After SPIDER assay, Pupylation site (K146) on WT CheAs was identified by LC-MS/MS analysis, by searching for an additional mass of ~243.09 Da, which represents the three C-terminal residues of Pup^E^, *i. e.*, GGE, that covalently binds to adjacent lysine by pupylation. **d,** SPIDER assays with Flag-tagged RutR and specific biotin-DNA. Motif-1: 5’-TTGACCATACGGTAAA-3’, Motif-2: 5’-ATGACCATGATTTCGT-3’. **e,** SPIDER assays with Flag-tagged Sox2 and specific biotin-RNA. Mutant: Sox2 binding RNA with multiple mutated sites. **f,** SPIDER assay with Biotin-Rapamycin and lysate of HEK293T cells, V5-tagged FKBP12 was overexpressed in these cells. **(d, e, f)** Quantitation of Relative abundance of the major band of SA-Pup^E^ ~ Prey based on western-blotting is shown at the right panel. Data are representative of one experiment with at least three independent biological replicates (mean and S.E.M. of n = 3), ** *p* < 0.01 and *** *p* < 0.001 (two-tailed unpaired t-test). **g,** Pupylation sites mapped on the crystal structure of FKBP12 (PDB: 1FKL).

### SPIDER proximity-labeling system for validating the protein-biomolecule interactions

Next, we examined whether SPIDER could efficiently validate the protein-biomolecule interactions. We chose the well-studied CheAs and CheZ interaction as a PPI model system (K_D_ ~350 nM (Extended Data Fig. 2a)^32^. We biotinylated CheZ, and as expected, CheAs could indeed be linked to SA-Pup^E^ through the binding with biotin-CheZ in a dose-dependent manner (Fig. 1b). Mass spectrometry analysis confirmed the success of the SPIDER reaction and showed SA-Pup^E^ was linked to CheAs at several lysine residues (Fig. 1c and Extended Data Fig. 2d,e). It is known that specific mutations of CheAs (L123A, L126A) reduce its affinity with CheZ^33^, which we also confirmed (Extended Data Fig. 2b,c).

To test the applicability of SPIDER for Protein-Nucleic acid interaction (PNI), we examined the interaction between RutR and DNA. RutR is the master regulator of genes involved in *E. coli* pyrimidine catabolism and is well known to bind to motif 5′-TTGACCANNTGGTCAA-3′^34^. As expected, we found that RutR was linked to SA-Pup^E^ through its binding to biotin-DNA and this linkage depends on the affinity of RuR to the DNA sequence (Fig. 1d)^34^. Excess non-biotinylated DNA effectively prevented the formation of SA-Pup^E^~RutR (Fig. 1d). MS analysis showed that the SA-Pup^E^ is linked to RutR at two lysine residues (Extended Data Fig. 2f,g).

As a test for the effectiveness of SPIDER with RNA-protein pairs, we examined Sox2 and the SARS-CoV-2 N protein. As a transcriptional regulator, Sox2 binds both DNA and RNA^35^, while N protein appears to exclusively bind RNA and regulates SARS-CoV-2 genome assembly^36^. We observed production of SA-Pup^E^~Sox2 upon the addition of specific biotin-RNA or biotin-DNA in a sequence-dependent manner (Fig. 1e and Extended Data Fig. 2h). A similar mobility-shift was also observed with the N protein interacting with RNA (Extended Data Fig. 2i). Further, we found that all of the pupylated lysines of both Sox2 and N protein are on their surfaces (Extended Data Fig. 2j to l). Interestingly, the pupylated lysines from Sox2~DNA (Extended Data Fig. 2j) and Sox2~RNA (Extended Data Fig. 2k) are different, and all of the modified lysines of the N protein (Extended Data Fig. 2l) are at the C-terminal region that rich of lysine residues, although the N-terminal of SARS-CoV-2 N is a RNA binding domain^36^. We expect that the sites on the proteins that are pupylated are dependent on their spatial relation with the nucleic acid binding region on the proteins. Thus, we conclude that Sox2 has different binding regions for DNA and RNA, which is consistent with previous study^35^.

To determine whether SPIDER could effectively detect Protein-Small molecule interaction (PSMI), we examined the interaction between rapamycin and FK-binding protein 12 (FKBP12). The binding of rapamycin to FKBP12 is a well-known mechanism to suppress the immune system^37^. We performed the SPIDER assay by incubating a Biotin-Rapamycin with the total lysate of FKBP12-overexpressed HEK293T cells. A significant production of SA-Pup^E^~FKBP12 was observed directly from the reaction mixture without enrichment (Fig. 1f), while the addition of excessive rapamycin effectively reduced the production of SA-Pup^E^~FKBP12 (Fig. 1f). Further, biotin agarose was used to enrich the SA-Pup^E^-linked proteins followed by stringent wash and MS identification. As expected, FKBP12 was significantly enriched and two pupylated lysine residues (K35 and K53) of FKBP12 were identified (Fig. 1g). Consistent with our observations of the spatial accessibility of SPIDER mediated pupylation on the target proteins, the identified pupylation sites K35 and K53 are indeed on the surface of a region adjacent to the rapamycin binding site (Fig. 1g).

Taken together, these results indicate that SPIDER is indeed highly effective for studying protein-biomolecule interactions.

### SPIDER-PPI to identify enzyme-substrate interactions

It is generally challenging to detect transient interactions, such as those between enzymes and their substrates, especially within a cellular milieu. To assess whether SPIDER could resolve these interactors, we identified the interactome of the *E. coli* protein deacetylase CobB^38^. This protein, the only member of the Sir2 family deacetylases in *E. coli*, is known to play a role in many different pathways^39,40^, although its complete range of interactors is still incompletely known. The SPIDER assay was carried out by incubating biotin-CobB with stable isotope labeled *E. coli* total lysate that was prepared under mild lysis condition (Fig. 2a and Extended Data Fig. 3a). A total of 84 interacting proteins was identified (Fig. 2f, Extended Data Fig. 3b and Supplementary Table 1). Notably, while half of the interacting proteins in our dataset were reported to be potential protein substrates of CobB^39,40^, we identified a variety of novel interactors (Fig. 2b and Supplementary Table 2). These interacting proteins are involved in 22 pathways, including gene expression that has also been found in previous studies ^39,40^ (Extended Data Fig. 3h). To validate these findings, we randomly selected eight proteins (Extended Data Fig. 3c) and examined them with biolayer interference (BLI), confirming the bindings for 5 of them, namely Protein-a (Fig. 2c), RraA, Protein-f, Protein-c and Protein-b (Extended Data Fig. 3d to g). Deacetylation assays clearly showed that Protein-b and Protein-c are acetylated in *E. coli*, and the acetylation could be efficiently removed by CobB (Fig. 2d,e). Protein-b (RNase R) is a 3’-5’ exoribonuclease whose activity is known to be regulated by acetylation^41,42^, whereas Protein-c is a critical component of the transcription initiation machinery. Our results thus imply that CobB plays a previously unappreciated role in regulating both of these processes. The CobB interactors identified here mainly cluster into four categories: namely, coenzyme metabolic process, gene expression, alpha-amino acid biosynthetic process and aromatic-amino acid family biosynthetic process (Fig. 2f). Collectively, these results show that SPIDER is indeed capable of identifying enzyme-substrate interactions within total cell lysate.

**Fig.2.**
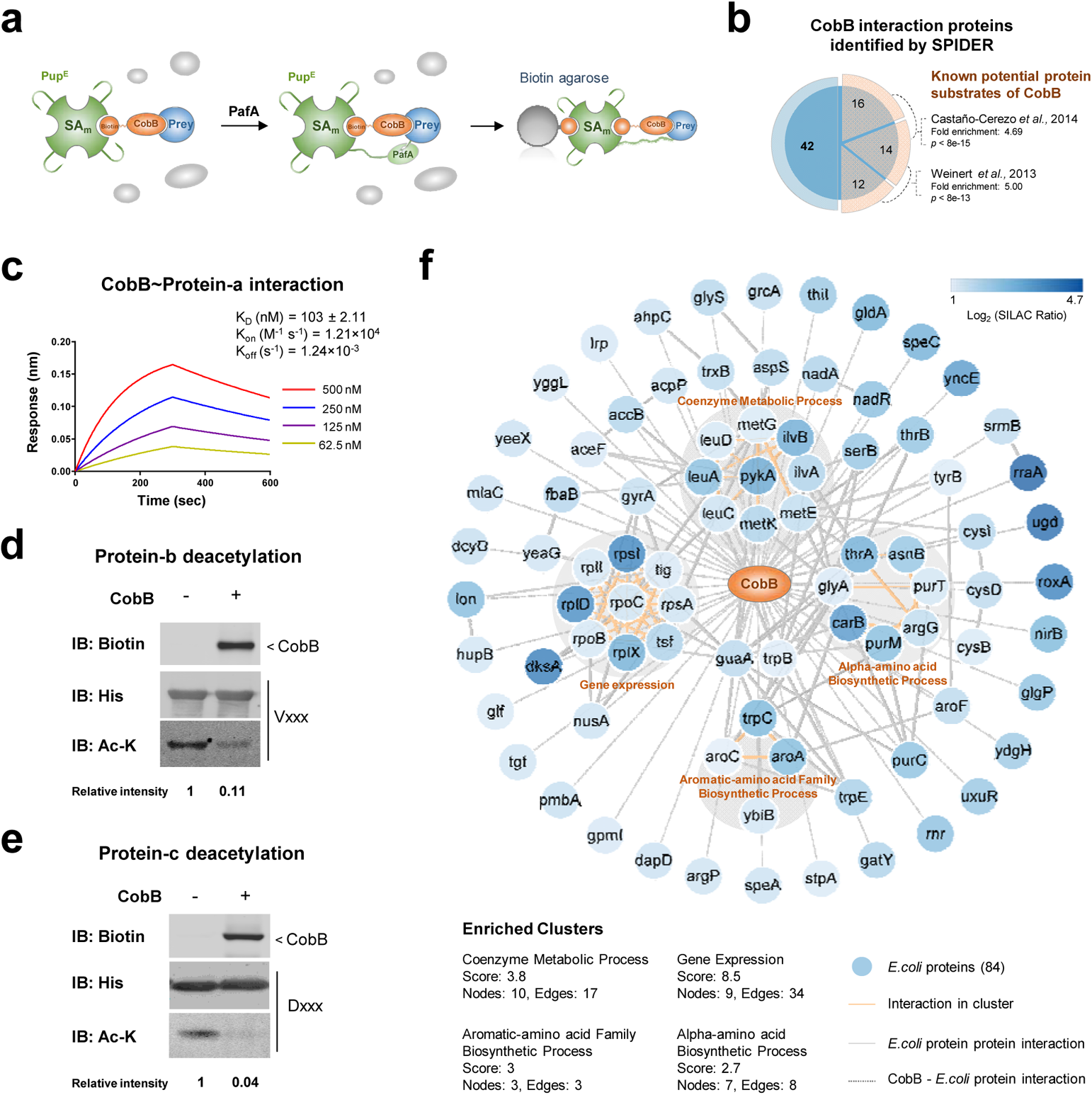
SPIDER-PPI to identify enzyme-substrate interactions. **a,** Schematic diagram of the application of SPIDER in capturing CobB interacting proteins. **b,** The 84 CobB interacting proteins identified by SPIDER. The known potential protein substrates of CobB that were identified in two studies (highlighted in orange). **c,** The newly identified binding proteins of CobB were validated by BLI, taking Protein-a as an example. **d, e,** Deacetylation assays of Protein-b **(d)** or Protein-c **(e)** with CobB. **f,** The interaction network of the CobB interacting proteins obtained from STRING with a confidence score > 0.7, by using the 84 interacting proteins as the input. The four top-ranked and tightly connected network clusters obtained with MCODE are color coded.

### SPIDER-PNI to identify *N^6^*-methyladenosine (m^6^A) binding proteins

Next, we asked whether SPIDER could identify modified nucleic acids binding proteins within a complex environment, such as the cellular milieu. To this end, we sought to identify *N*^6^-methyladenosine (m^6^A) binding proteins. m^6^A is the most prevalent modification on eukaryotic mRNA and is interpreted by its readers, such as YTH domain-containing proteins, to regulate the fate of mRNAs^43,44^. We carried out the SPIDER assay by incubating biotin-ssRNAs (with and without m^6^A modification) with the total lysate of YTDHFs (YTHDF1, YTHDF2 and YTHDF3)-overexpressed HEK293T cells. The total lysate was prepared under mild lysis conditions (Fig. 3a). As expected, gel-shift electrophoresis assays confirmed the formation of SA-Pup^E^~YTHDFs directly from the reaction mixture without enrichment, which could be reduced by excessive non-biotinylated m^6^A (Fig. 3b and Extended Data Fig. 4a,b). Further, using MS to identify proteins linked to SA-Pup^E^, we found that the YTDHFs scored in the top 3 among the 20 identified proteins. Two known m^6^A readers, IGF2BP2^43^ and YTHDC1^45^, were also identified among the top interactors (Fig. 3c and Supplementary Table 3). Gene ontology analysis showed that more than half of the identified proteins are well connected to the m^6^A reader signaling pathway and clustered into two categories: regulation of mRNA metabolic process and fidelity maintenance of DNA replication (Fig. 3d and Extended Data Fig. 4e). Further, when we analyzed the pupylation sites on YTDHFs by MS, we found that the modified lysine sites were highly conserved among the YTDHFs and all are located on the protein surface adjacent to the m^6^A binding region (Fig. 3e,f and Extended Data Fig. 4c,d). Collectively, these results show that SPIDER is indeed suitable for identifying PNIs of modified nucleic acids in a cellular milieu.

**Fig.3.**
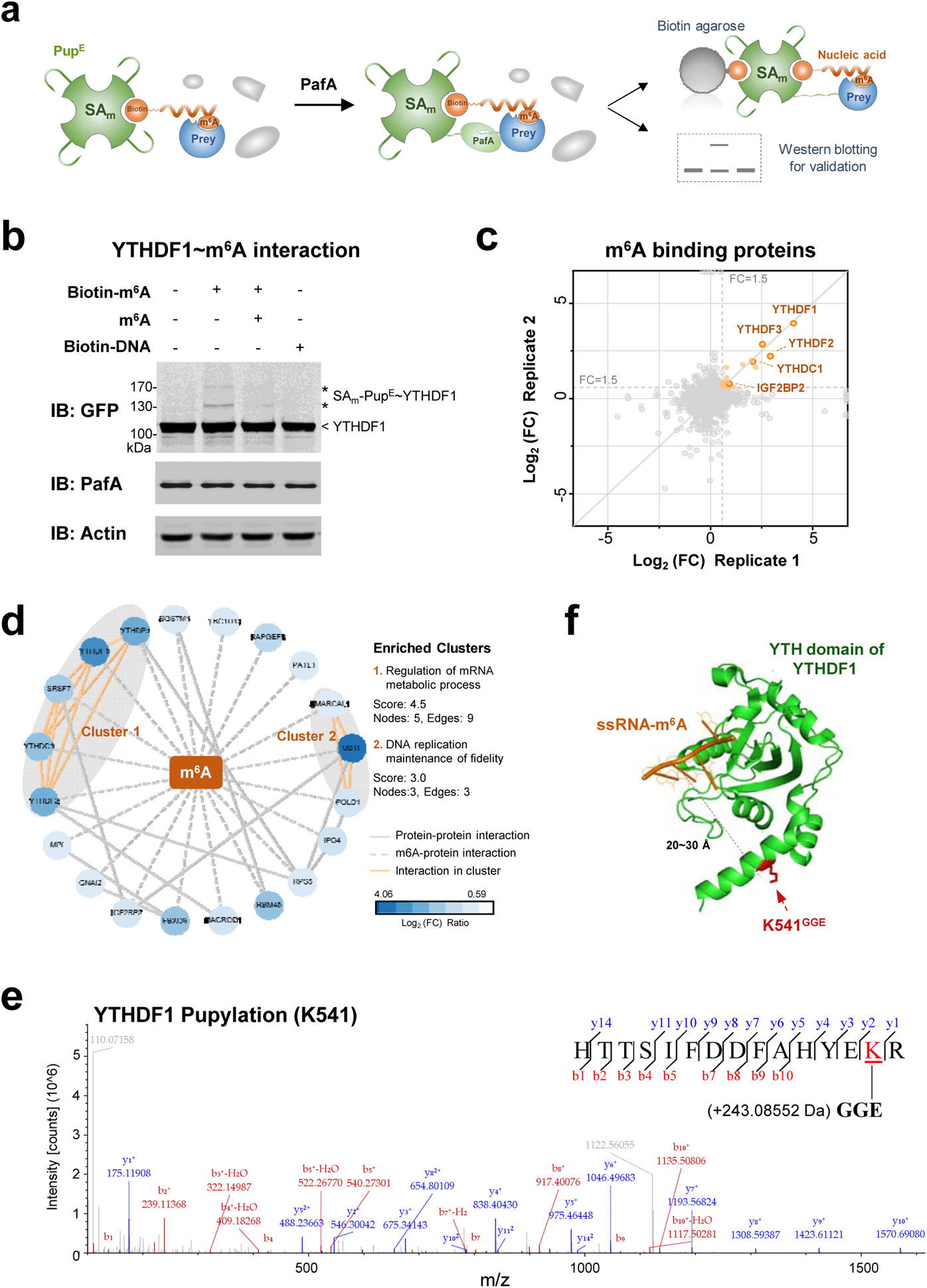
SPIDER-PNI to identify N^6^-methyladenosine (m^6^A) binding. **a,** Schematic diagram of SPIDER in capturing m^6^A interacting proteins. **b,** SPIDER assays with the cell lysates of YTHDF1-overexpressed HEK293T cells and biotin-m^6^A. **c,** Scatter plots show comparison of fold change of two replicate experiments of SPIDER~Biotin-m^6^A. The y axis represents the log2 fold change of label-free protein quantification (SPIDER~Biotin-m^6^A / SPIDER~Biotin-ssRNA). Significantly enriched proteins are shown as orange dots. FC: fold change. **d,** The interaction network of the m^6^A-interacting proteins obtained from STRING with a confidence score > 0.4, by using the 20 interacting proteins identified by SPIDER as input. The two top-ranked and tightly connected network clusters obtained with MCODE are grey color coded. **e,** Pupylation site (K541) on YTHDF1 after SPIDER assay, was identified by LC-MS/MS analysis. **f,** Pupylated site mapped on crystal structure of YTHDF1 YTH domain in complex with 5-mer m^6^A RNA (PDB: 4RCJ). The pupylated lysines are highlighted in red.

### SPIDER-PSMI to capture the interacting proteins of lenalidomide

Next, we asked whether SPIDER could capture PSMIs in a complex environment. To answer this, we employed SPIDER to identify the targets of a widely used medicine, lenalidomide^46^. Lenalidomide is well-known for its effectiveness to treat hematological malignancies via its interaction with the E3 ligase DDB1-CRBN complex^46^. We expected that either CRBN, or DDB1 could be captured in the SPIDER assay (Fig. 4a). We carried out the SPIDER assay by incubating Biotin-Lenalidomide or a control of biotin only with HEK293T total lysate, followed by the use of biotin agarose coupled with MS to identify the interacting proteins (Fig. 4a and Extended Data Fig. 5a). As expected, the DDB1-CRBN complex was highly enriched by Biotin-Lenalidomide (Fig. 4b and Supplementary Table 4). We found that the pupylated lysines were all located on the protein surface within close proximity to the binding interface, reflecting sites of greatest access to the SPIDER reaction (Fig. 4c,d and Extended Data Fig. 5b,c). In addition, besides CRBN and DDB1, SPIDER also identified other interacting proteins for lenalidomide (Fig. 4b, 4e and Supplementary Table 4). Gene ontology (GO) analysis showed that more than half of the identified proteins are well connected to the CRBN-DDB1 signaling pathway and clustered into the following categories: nucleotide-excision repair, mitochondrial translation and Golgi vesicle transport (Fig. 4e and Extended Data Fig. 5d). These binding proteins could serve as a resource to explore other therapeutic purposes of lenalidomide or possibly explain the known side-effects^47^. Collectively, these results show that SPIDER can identify target proteins of small molecular drugs within a complex environment such as the cellular milieu.

**Fig.4.**
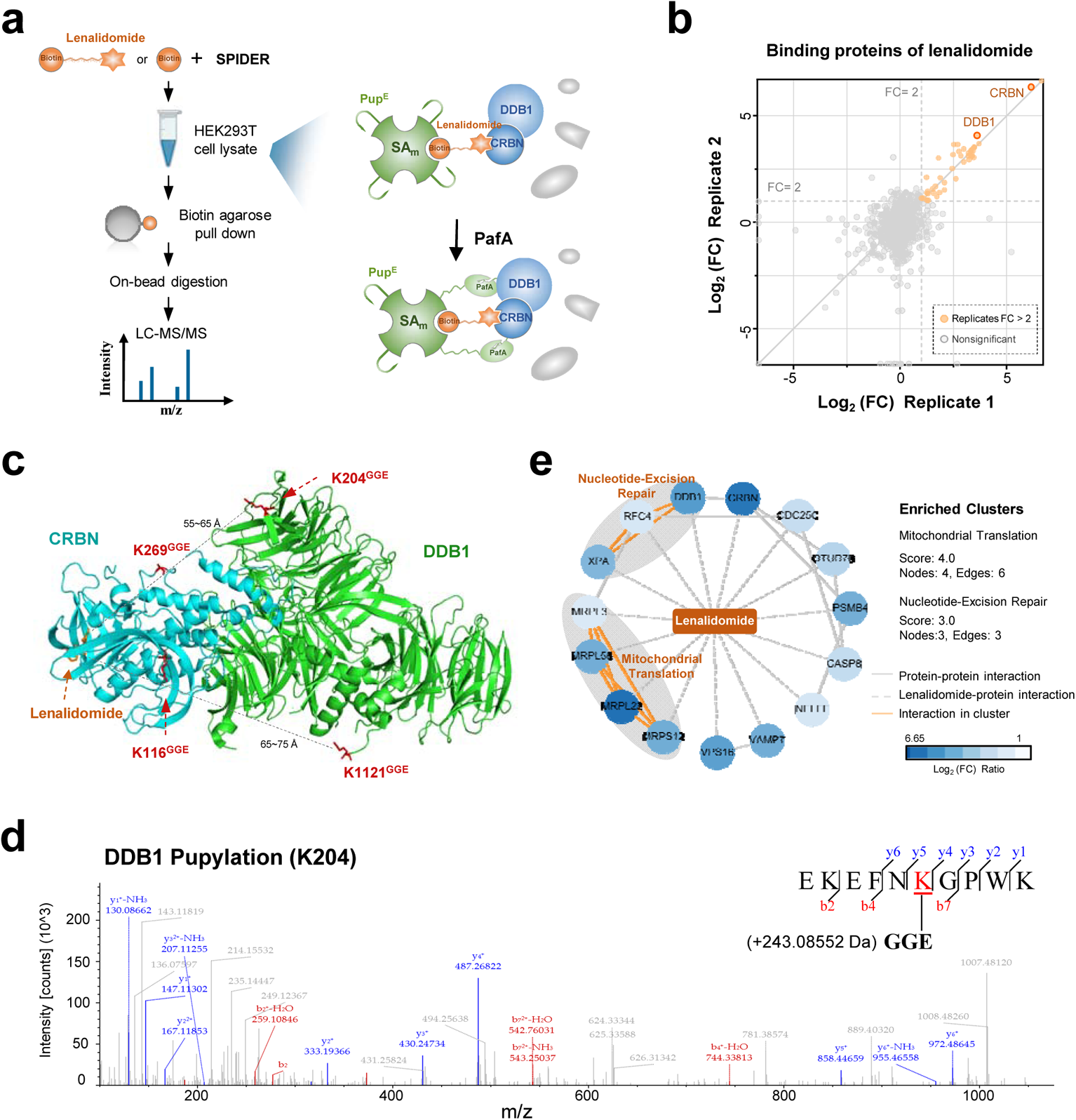
SPIDER-PSMI to capture the interacting proteins of lenalidomide. **a,** The workflow of SPIDER assays for capturing the binding proteins of biotinylated lenalidomide. Cartoon of SPIDER in capturing the Lenalidomide~CRBN-DDB1 complex is shown at the right panel. **b,** Scatter plots show comparison of fold change (FC) levels of two replicate experiments of SPIDER~Biotin-Lenalidomide. The y axis represents the log_2_(FC) of SPIDER~Biotin-Lenalidomide / SPIDER~Biotin). Significantly enriched proteins are shown as orange dots. **c,** Pupylation sites on the crystal structure of Lenalidomide~CRBN-DDB1 complex (PDB: 4CI2). Pupylated lysines are highlighted in red. **d,** The pupylation site (K204) on DDB1 after SPIDER assay identified by LC-MS/MS analysis. **e,** The interaction network of the lenalidomide-interacting proteins obtained from STRING with a confidence score > 0.4, by using the 20 interacting proteins identified by SPIDER as input. The two top-ranked and tightly connected network clusters obtained with MCODE are grey color coded.

### SPIDER to identify SARS-CoV-2 Omicron variant receptors on cell surface

In comparison to protein-biomolecule interactions among soluble proteins, the study of protein-biomolecule interactions that involve membrane proteins like ligand-receptor interaction is considerably more challenging due to the difficulty to isolate membrane proteins while maintaining their conformations and activities.

The SARS-CoV-2 Omicron variant (B.1.1.529) is of great concern. Compared to the Prototype, Omicron carries more than 32 mutations on the Spike protein. Recent studies showed Omicron has extensive immune escape of neutralizing antibodies, convalescent and vaccinated sera^48–50^. Omicron variant infects more easily than the Delta variant and the Prototype in human bronchus, but with less severe lung infection^51^. Unveiling the molecular mechanism of the high transmissibility of Omicron is essential for developing protective strategies. Mutations in the receptor-binding domain (RBD) of Spike protein may affect tropism of SARS-CoV-2 and consequently pathogenesis. There are 15 mutations in the Omicron RBD, however, the binding affinity difference to hACE2 between the Omicron RBD and the Prototype is still controversial^49,52^. Thus, it is highly possible that other critical host receptors and/or co-receptors might promote the binding and entry of SARS-CoV-2 into human bronchus but not lung.

To test the capability of SPIDER for identifying cell-surface receptor (Fig. 5a). We first examined the interaction between the SARS-CoV-2 S1 protein and its well-known receptor, ACE2. SPIDER reactions were carried out directly on cells in petri dish. We observed no change of the cell mortality before and after the SPIDER reaction (Extended Data Fig. 6a). The binding between SARS-CoV-2 S1 and ACE2 was confirmed by showing the mobility shift of ACE2-SA-Pup^E^ (Fig. 5b).

**Fig.5.**
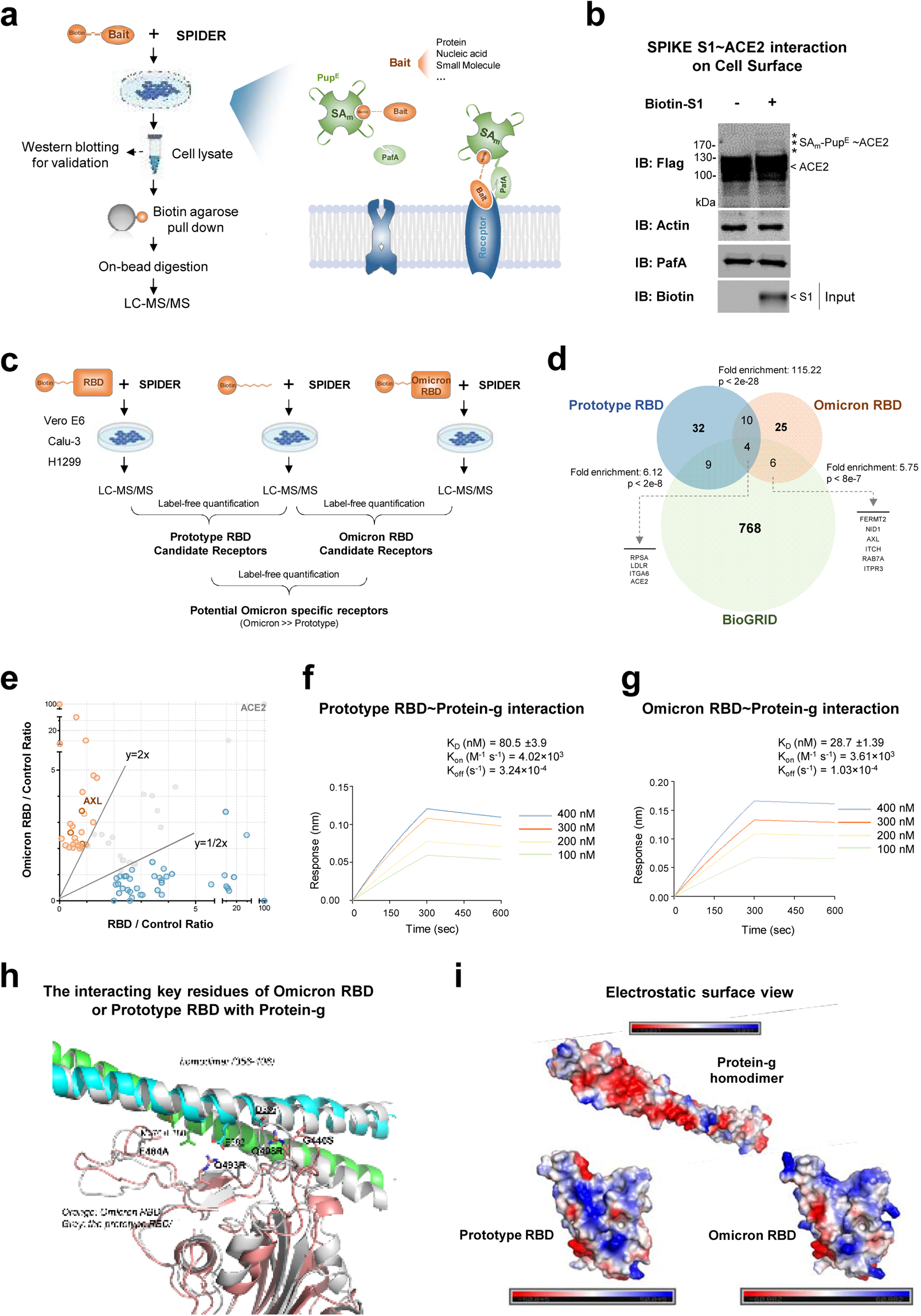
SPIDER to identify SARS-CoV-2 receptor on cell surface. **a,** The workflow of SPIDER assays for the identification of biotinylated bait binding proteins on cell surface. **b,** SPIDER assay with the biotin-S1 and lysate of HEK293T cells, Flag-tagged ACE2 was overexpressed in these cells. The asterisks represent the bands of SA-Pup^E^ ~ACE2. **c,** The workflow of SPIDER assays for the identification of biotinylated RBD and Omicron RBD binding proteins on cell surface. The prey protein/s is identified by mass spectrometry. **d,** The Prototype RBD and the Omicron RBD interactors identified by SPIDER were compared with known interactors in the BioGRID database. **e,** Scatter plots represent the ratio of label-free protein quantification of the two experiments, *i. e*., SPIDER~Biotin-Omicron RBD and SPIDER~Biotin-RBD. Significantly enriched proteins for Omicron RBD and Prototype RBD are shown as orange and blue dots, respectively. **f,g,** Binding affinity of the Prototype RBD~Protein-g **(f)** and Omicron RBD~Protein-g **(g)**. **h,** Interaction patches in predicted complex structures of the Protein-g (Orange)-Omicron RBD and the Prototype RBD. **i,** Electrostatic surface view of Protein-g, Prototype SARS-CoV-2 RBD and Omicron RBD.

Next, we coupled SPIDER with MS to identify the Prototype RBD and Omicron RBD binding proteins on the surface of H1299, Calu-3 and Vero E6 (Fig. 5c and Extended Data Fig. 6b). H1299 and Calu-3 cells represent low and high expression of ACE2, respectively. We identified 55 and 45 interactors for Prototype RBD and Omicron RBD, respectively (Fig. 5d and Supplementary Table 8), including several of the known receptor/co-receptors, *e. g.*, ACE2, AXL^53^ and Protein-g. To further confirm the ligand-receptor binding, based on the MS data, the pupylated lysine sites were identified (Extended Data Fig. 6c,d). The pupylated site is on the surface of ACE2 that is accessible to the SPIDER reaction (Extended Data Fig. 6e). After these identified RBD interactors and the interactors in the BioGRID database were compared, we found 4 interactors, *i. e*., ACE2, ITGA6, RPSA and LDLR, are overlapped among all these three groups (Fig. 5d and Supplementary Table 6).

To further explore the functional roles of the potential receptors, we studied the high confidence interactors of Omicron RBD in regarding to their protein interaction network, protein level, mRNA level and potential drugs. We first analyzed the protein interaction network of Prototype RBD and Omicron RBD (Extended Data Fig. 7a,b), we identified 3 major cell processes of the interactors of Omicron RBD, *e. g.*, focal adhesion. We found 29 proteins were significantly enriched in Omicron RBD vs. Prototype RBD (Fig. 5e and Supplementary Table 7). Among these proteins, Protein-g is highly conserved within species (Extended Data Fig. 7c). It is a major component of class-III intermediate filaments and also known to localize on the extra-cellular surface of the plasma membrane, as a co-receptor participating in SARS-CoV-2 infection. We performed Biolayer interferometry (BLI) assays and confirmed the bindings of Prototype RBD, Omicron RBD and Prototype S1 to Protein-g (Fig. 5f,g and Extended Data Fig. 7d). Protein-g was confirmed to selectively bind to the Omicron RBD with an approximately 3 folds higher affinity than that of the Prototype RBD.

To analyze the protein level and the mRNA level of the Omicron RBD specific enriched proteins (Supplementary Table 7) among different tissues, especially among nasopharynx, bronchus and lung, we adopted data from the Human Protein Atlas (HPA)^54^ and a published single-cell mRNA sequencing dataset of 232,905 single cells from major adult human organs (all from non-SARS-CoV-2-infected samples)^55^. We found that many of the interactors including Protein-g were higher in trachea than lung, both at protein and mRNA levels (Extended Data Fig. 7e,f and Supplementary Table 8). These results may at least partially explain the significant higher proliferation rate of the Omicron variant at trachea than in lung^56^.

To unveil the mechanism underlying the binding between the Omicron RBD and Protein-g, we performed docking analysis. The crystal structure of Protein-g C2 domain (PDBID, 1GK4) was used as the input structure for the docking. Based on the fact that the Protein-g binding affinities of the Omicron RBD and the Prototype were different, only the docking solutions with at least 4 mutations of the Omicron RBD at the interface of RBD-Protein-g were filtered out, and the best one was selected for refinement and further analysis. The binding structure of Protein-g and the Prototype RBD was built using this model as a template. Extended Data Figure 8a and 8b shows the refined complex models of human Protein-g and the two types of RBD. Molecular dynamics simulations of these two predicted RBD-Protein-g complex models were performed for the model relaxation and binding free energy estimation. Using the MM/GBSA method based on 18 ns trajectories of each model, the calculated binding free energies of the Omicron RBD and the Prototype RBD to Protein-g are −49.7 ± 9.0 kcal/mol and −15.7 ± 8.7 kcal/mol, respectively. Thus, Omicron RBD-Protein-g shows a significant stronger binding than Prototype RBD-Protein-g, which correlated well to the measured binding affinities. Figure 6h shows the Omicron RBD-Protein-g and the Prototype RBD-Protein-g complexes. For the Omicron RBD, a E484A mutation is more favorable for interacting with the hydrophobic residues of Protein-g M376 and L380; Q493R and Q498R mutations helps to form strong electrostatic interaction with Protein-g E382 and D385. Moreover, Omicron RBD G446S may form additional hydrogen-bonding with Protein-g (Extended Data Fig. 8c highlighted the 4 mutated residues). We plotted the electrostatic surface of Omicron RBD, the Prototype RBD and Protein-g (Fig. 5i). It is clear that Protein-g has a negative-charged surface, and the Omicron RBD Q493R and Q498R makes it more suitable for the electrostatic complement.

Taken together, the Omicron RBD enriched potential receptors could be used to explain the differences between the Omicron variant and the Prototype strain. Some of the candidates could be further investigated as drug targets to specifically combat the Omicron variant. These results show that SPIDER can efficiently identify membrane-localized receptors.

## Discussion

Conceptually, SPIDER is based on a mechanism whereby the covalent linkage happens when the substrate (Pup^E^) is proximal to protein/s. To our knowledge, SPIDER is the only substrate-based proximity labeling system.

There are several notable advantages of SPIDER. Firstly, SPIDER is generally applicable for any biological molecules, with the only prerequisite being the biotinylation of the molecule of interest, and biotinylation could be easily achieved through synthesis or labeling *in vitro* or *in vivo*. And the small biotin modification is less likely to interfere with protein-biomolecule interactions than other methods, such as tri-functional affinity probes or enzyme-fused bait proximity labeling. In addition, specific modifications, *e.g*., m^6^A, could also be readily incorporated into the bait of interest, which enable SPIDER for identifying of modification-mediated protein-biomolecule interaction. Secondly, SPIDER is relatively simple to perform and highly reproducible. In a typical SPIDER assay, the interaction between the bait and the prey protein is transformed into a covalent linkage between the prey protein and SA-Pup^E^ through enzymatic proximity pupylation at mild condition. After incubation, without concern about the degradation of the bait, especially RNA, and the destruction of protein-biomolecule interaction, especially ligand-receptor on cell surface, the reaction could be subjected to extremely stringent washes without losing the specific bindings. In addition, because of the covalent linkage, the results of SPIDER could be easily visualized on a gel by simply monitoring a mobility shift of the SA-protein conjugate. This feature is very suitable for fast *in vitro* validation of protein-biomolecule interactions. Thirdly, due to the spatial controllability of SPIDER, we can roughly predict the bait binding position on the protein through the analysis of pupylation sites. By combining this data with protein structure, and bioinformatic tools, such as docking, we expect to be able to pinpoint the key residues involved in protein-biomolecule interaction and in-depth mechanistic understanding. Lastly, the key reagents of SPIDER (PafA and SA-Pup^E^) could be prepared in large quantity, which allows a batch SPIDER trials to investigate a large number of different baits interactor at one time.

Nonetheless, there are a few limitations of SPIDER. Firstly, in its current form, SPIDER is basically an *in vitro* system, although as demonstrated here, ligand-receptor interactions on an intact cell surface can be identified. The concept of substrate-based proximity activity could also be applied to establish an *in vivo* system by simply fusing the 27~64 amino acids of pup^57^ to a bait protein or fusing pup to dCas proteins, which binds the target nucleic acid through sgRNA. Secondly, similar to enzyme-based proximity labeling methods, unwanted self-labeling is inevitable. To minimize the level of self-labeling, we mutated all the surface-exposed lysine residues on the three key proteins (Pup^E^, PafA and SA) and still maintained comparable activities. Third, non-specific binding is another concern. This limitation can be largely overcome by adopting SILAC (Stable Isotope Labeling of Amino Acids) strategy.

By applying SPIDER, in a very short period of time, we identified 29 Omicron RBD enriched interactors on cell surface, including Protein-g. These interactors could serve as lead candidates for developing Omicron variant specific therapeutics. The binding of Protein-g to the Omicron RBD is approximately 3 folds higher than that of the Prototype RBD. Docking and Structure analysis revealed that, comparing with Prototype RBD, the binding surface of Omicron RBD has a larger positive charge region. Protein-g is a strongly negative charged polyelectrolyte, usually mediate interactions with other molecules through the negative charge. The larger positive charge region of Omicron RBD may account for the higher affinity with Protein-g. In addition, the trachea has higher level cell-surface Protein-g. Because of these observations, we reason that during infection, more SARS-CoV-2 Omicron variant virions could be trapped on the trachea surface than that of the lung, resulting in much higher proliferation rate of Omicron in human bronchus than lung^56^.

To conclude, SPIDER is the first substrate-based proximity labeling system. Because of the nature of enzymatically transform non-covalent interaction to covalent interaction, it enables efficient and specific identification and validation of protein-biomolecule interactions, especially for weak, transient and membrane-localized interactions, as long as the biomolecule could be biotinylated, either *in vitro* or *in vivo*.

## Acknowledgments

We thank Dr. Heng Zhu for his long-term guidance.

We thank Prof. Pilong Li of School of Life Sciences of Tsinghua University for kindly providing the YTHDFs family plasmids.

We thank Prof. Xichen Bao of Guangzhou Institutes of Biomedicine and Health of Chinese Academy of Sciences for kindly providing the Sox2 plasmid.

We thank Prof. Xiaodong Zhao of Shanghai Jiao Tong University and Prof. Jian Yang of Tongji University for critical comments.

## Author contributions

Conceptualization: S.C.T, H.W.J.

Investigation: H.W.J., H.C., Y.X.Z., X.N.W., Q.F.M., X.J., J.Z., C.S.Z., and J.F.P.

Resources: Z.W.X., Z.Q.C., L.W., W.S.K., K.Z., M.L.M., H.N.Z., S.J.G., J.B.X., J.L.H., Z.Y.L., W.X.N., F.J.W., T.W., W.L., Y.J.D., R.N.W., Y.J.D., Y.Q. and J.J.D.

Writing – original draft: S.C.T, J.F.P. and H.W.J.

Writing – review & editing: D.M.C.

## Data and materials availability

The Mass spectrometry proteomics data have been deposited to the ProteomeXchange Consortium (http://proteomecentral.proteomexchange.org) via the iProX^1^ partner repositor with dataset identifiers as follows:

PXD026509: CheAs pupylation site PXD026509: CheAs pupylation sites identified by SPIDER assay

PXD026527: Intra-molecule pupylation of GFP-Pup^E^ identified by mass spectrometry

PXD026511: Pupylation sites on protein FKBP12

PXD026478: Pupylation sites on protein GFP-Pup^E^

PXD026514: CobB interacting proteins identified by SPIDER assay

PXD026516: Pupylation sites on protein RutR;

PXD026517: Pupylation sites on protein Sox2;

PXD026518: Pupylation sites on SARS-CoV-2 N;

PXD031035: SARS-CoV-2 receptor on cell surface identified by SPIDER assay.

PXD026523: Lenalidomide binding proteins identified by SPIDER assay

Additional data related to this paper may be requested from the authors.

## Methods

### Plasmid construction and cloning

The protein sequences were downloaded from GenBank. The corresponding DNA sequences were codon-optimized and synthesized by Sangon Biotech (China). The synthesized genes were cloned into pET28a or pGEX-4T-1 for prokaryotic expression or pcDNA3.1 for eukaryotic expression. PafA was constructed based on wild-type PafA using a QuickChange®. The DNA sequences of Streptavidin and Pup were fused together through PCR. The PCR product (SA-Pup) was cloned into pET28a. Proteins were expressed in *E. coli* BL21(DE3). N- or C-terminal His and Avi (GLNDIFEAQKIEWHE) tagged proteins of interest (POI) were cloned into pET32a, the pET32a plasmids were co-transformed with pET28a carrying BirA into *E. coli* BL21(DE3). All biotin-DNA were synthesized in Sangon Biotech (China) and all biotin-RNA were synthesized in Genscript (China).

### Protein expression and purification

*E. coli* BL21, carrying the expression plasmid, was cultured in LB medium. Protein expression was induced with isopropyl-β-D-thiogalactoside (IPTG). For the purification of 6xHis-tagged proteins, cell pellets were re-suspended in lysis buffer, then lysed by the high-pressure cell cracker (Union-biotech, China). Cell lysates were centrifuged at 12,000 rpm for 20 mins at 4. Supernatants were collected and purified with Ni^2+^ Sepharose beads (Smart-lifesciences, China) or Ni-NTA Agarose (QIAGEN, Germany), then washed with lysis buffer (For the purification of biotinylated protein, biotin is added to the wash buffer) and eluted with elution buffer. For the purification of GST-tagged proteins, cells were harvested and lysed by the high-pressure cell cracker in lysis buffer. After centrifugation, the supernatant was incubated with GST-Sepharose beads (Smart-lifesciences, China) or Glutathione Sepharose 4 Fast Flow (Cytiva, USA). The target proteins were washed with lysis buffer twice and eluted with elution buffer.

For recover soluble SA-Pup^E^ for inclusion body, the insoluble fraction cultured from 200 mL LB medium was washed four times. The inclusion body was dissolved in 8 M urea and dialyzed against 0.8 M urea and then dialyzed against 50 mM Tris-HCl buffer for another 24 h. The dialysate was centrifuged and concentrated in a stirred ultrafiltration cell (Amicon, USA).

### Cell culture and transfection

HEK293T cells, Calu-3 cells, Vero E6 cells, and H1299 cells were cultured in DMEM medium (Corning, USA) by adding 10% fetal bovine serum (FBS) (Excel, China) and 1% penicillin-streptomycin (Invitrogen, USA) at 37 in a 5% CO_2_ incubator. During passaging, cells were digested with 0.25% trypsin. Before transfection, cells were passaged by 1:3~1:5 and cultured for 12-16 h. The cells were transfected using Lipofectamine™ 2000 transfection reagent (Invitrogen, USA) according to the manufacturer’s instruction. Briefly, in a 10-cm petri dish, 10 μg plasmid and 25 μL Lipofectamine™ 2000 were added. After transfection, cells were cultured for 36-48 h.

For SILAC labeling of HEK293T cells, 50 mg of ^13^C^6^ L-Lys and 50 mg of ^13^C_6_ ^15^N_4_ L-Arg were added to 0.5 liter of −Lys/−Arg SILAC DMEM (Thermo, USA) supplemented with 10% FBS and 1% penicillin-streptomycin to generate “Heavy” medium. “Light” medium cells were cultured using DMEM supplemented with 10% FBS, and 1% penicillin-streptomycin.

### Cell lysate preparation

For adherent cells, cells were washed once in PBS buffer. 1 mL M-PER Reagent (Thermo Scientific, USA) with 0.5 mM PMSF was added to a 10 cm petri dish, followed by 5 min gentle shake to obtain the cell lysate. For suspension cells, cells were counted using a cell counting chamber. Cell suspension was centrifuged at 800 rpm for 3 min. Cell pellet was washed with PBS and centrifuged at 800 rpm for 3 min. Supernatant was carefully removed. 1 mL M-PER Reagent with 0.5 mM PMSF was added to 1 mL/10^7^ cells, and vortexed briefly to obtain a homogeneous cell suspension. Cell suspension was incubated for 40 minutes at 4°C, with a 10-level speed (the highest speed) vortex every 10 minutes. Lysate was collected and transferred to a microcentrifuge tube, and centrifuged at 16,000 × *g* for 5-10 min to pellet the cell debris. Supernatant was transferred to a new tube and stored at −80°C for future use.

### SPIDER assay for PPIs validation

The reaction is composed of PafA, biotin-POI (protein of interest), purified prey protein, SA-Pup^E^ and ATP in reaction buffer. The reaction buffer includes 150 mM NaCl, 50 mM Tris-HCl, pH 7.5, 20 mM MgCl_2_, and 10% glycerol. After gentle mixing, the reaction is carried out for 4 h.

### SPIDER assay for PNIs validation

The reaction is composed of PafA, biotin-nucleic acid, purified prey protein, poly d(I-C), SA-Pup^E^ and ATP in reaction buffer. Unlabeled nucleic acid as a competitive probe was added to the reaction mixture when necessary. The reaction buffer includes 150 mM NaCl, 50 mM Tris-HCl, pH 7.5, 20 mM MgCl_2_, and 10% glycerol. After gentle mixing, the reaction is carried out for 4 h.

### SPIDER assay for PPIs screening in cell lysate

For SPIDER reaction, there are cell lysate, biotinylated POI or biotin control and PMSF, the reaction buffer is composed of 50 mM Tris-HCl pH 7.5, 150 mM NaCl, 20 mM MgCl_2_, 10%(v/v) glycerol. The reaction is carried out on a rotating wheel at room temperature, followed by the addition of ATP, SA-Pup and PafA, and incubation on a rotating wheel for 4~6 h. Save 10~30 μL sample for western blot analysis. Urea powder was added to the sample to a final concentration of 8 M. Biotin-agarose (Sigma, USA) was added to the reaction and incubated overnight to capture the pupylation linked interacting proteins. The biotin-agarose was collected and stored at −80°C for Mass spectrometry analysis.

For SILAC cell lysate, SPIDER assays were performed as described above with the “Heavy” (heavy isotope labeled) reaction containing biotinylated POI and the “Light” reaction containing biotin control. After incubation, the lysate was denatured by the addition of urea to a final concentration of 8M. The “Heavy” and “Light” reactions where then combined, and the mixture was incubated with biotin-agarose (Sigma, USA) overnight to capture the pupylation linked interacting proteins. The biotin-agarose was collected and stored at −80°C for Mass spectrometry analysis.

### SPIDER assay for PNIs screening in cell lysate

For SPIDER reaction, there are cell lysate (approximately 2~3 × 10^7^ cells), m^6^A ssRNA or control for m^6^A binding protein assay or Oligo (dT) for mRNA interactome assay, poly d(I-C) and 0.5 mM PMSF, the reaction buffer is composed of 50 mM Tris-HCl pH 7.5, 150 mM NaCl, 20 mM MgCl_2_, 10% (v/v) glycerol. The reaction is carried out on a rotating wheel at room temperature, followed by the addition of ATP, SA-Pup^E^ and PafA, and incubation on a rotating wheel for 4~6 h. Save10~30 μL sample for western blot analysis. Urea powder was added to the sample to a final concentration of 8 M and incubated for 2 min. Biotin-agarose (Sigma, USA) was added to the reaction and incubated overnight to capture the pupylation linked interacting proteins. The biotin-agarose was collected and stored at −80°C for Mass spectrometry analysis.

### SPIDER assay for PSMIs screenings in cell lysate

In a reaction, there are cell lysate, biotin-small molecule or biotin control, and 0.5 mM PMSF. The reaction buffer is composed of 50 mM Tris-HCl pH 7.5, 150 mM NaCl, 20 mM MgCl_2_, 10%(v/v) glycerol. The reaction was carried out on a rotating wheel at room temperature, followed by the addition of ATP, SA-Pup^E^ and PafA, and incubation on a rotating wheel for 4-6 h. Save 10-30 μL sample for western blotting analysis. Urea powder was added to the sample to a final concentration of 8 M, and incubated for 2 min. Biotin-agarose (Sigma, USA) was added to the reaction and incubated overnight to capture the pupylation linked interacting proteins. The biotin-agarose was collected and stored at −80°C for Mass spectrometry analysis.

### SPIDER reaction on cell surface

The adherent cells (HEK293T, Vero E6, H1299, Calu-3) were cultured in petri dish. After the removal of the medium, the cells were washed gently with the reaction buffer. biotinylated POI, SPIKE Prototype RBD or Omicron RBD (ACROBiosystems, China), nucleic acid and Biotin-Hexapeptide (synthesized by Sangon Biotech, China), or the control (free biotin only) were incubated with the cells for 15 minutes in the reaction buffer. The reaction buffer contained 50 mM Tris-HCl pH 7.5, 150 mM NaCl, 20 mM MgCl_2_, 10%(v/v) glycerol. Then, SA-Pup^E^, ATP and PafA were added to the reaction system and incubated for 2 h. The reaction system was removed, the cells were lysed by Radioimmunoprecipitation assay buffer (RIPA) (Beyotime Biotechnology, China) for 40 minutes. Save 10-30 μL sample for western blot analysis. Then Urea powder was added to the reaction to a final concentration of 8 M. The lysate was incubated on a rotating wheel at room temperature until Urea dissolved. After centrifugation of cell lysate, biotin agarose (Sigma, USA) was added to the supernatant and incubated overnight to capture the pupylation linked interacting proteins on cell surface. The biotin-agarose was collected and stored at −80°C for Mass spectrometry analysis.

### Mass spectrometry analysis of intact protein

The mass of GFP-Pup proteins from experimental and control groups were determined by Thermo Exactive Plus EMR mass spectrometry coupled to Agilent 1100 HPLC system. The proteins were first buffer exchanged into 0.1% formic acid with ultrafiltration. Then 0.5 μg proteins were loaded onto a 5 cm × 200 μm i.d. trap column (C5, 5 μm, 300 Å, Phenomenex) and separated by a 15 cm × 150 μm i.d. analytical column (C5, 5 μm, 300 Å, Phenomenex) with isocratic elution (50% acetonitrile 0.1 % formic acid). The mass spectra were collected with a resolution of 17,500 in Orbitrap analyzer. The EMR mode was enabled. The AGC (automatic gain control) was set at 1×10^6^ with an injection time of 100 ms. Finally, the mass spectra were deconvoluted with Thermo Protein Deconvolution 4.0 to get the exact mass of the proteins.

### Mass spectrometry and data analysis

Proteins enriched by biotin-agarose were reduced by adding DTT to a final concentration of 10 mM and incubated at 37 for 1 h. Subsequently, alkylation was performed by adding iodoacetamide to a final concentration of 25 mM and incubated in dark for 20 min. Proteins on the biotin-agarose were then digested with trypsin (1:30 protein-to-enzyme ratio) at 37 overnight. The biotin-agarose was rinsed twice with 200 μL 50 mM NH_4_HCO_3_. All the supernatant including the resulting peptides were collected and desalted using MonoSpin C18 care desalting column (GL Science, Japan) according to the instruction.

The tryptic peptide digests of the proteins were analyzed with an EASY-nL 1200 system coupled online to a Q Exactive plus Mass spectrometer (Thermo Scientific, USA). The peptide sequences were determined by searching MS/MS spectra against the Protein database using the Proteome Discoverer (version 2.4, Thermo Scientific, USA) software suite with a precursor ion mass tolerance of 10 ppm and fragment ion mass tolerance of 0.02 Da. Carbamidomethyl (C) was set as the fixed modification, oxidation (M), GGE (K) and deamidated (NQ) were set as the variable modification. The search results were automatically processed at the FDR 1% at both the protein and peptide. The unique peptide included in the protein group were used for quantification. All proteins were identified with unique peptide more than one. Label-free quantification was used to quantify the difference in protein abundance between different samples^2^.

For SILAC MS data analysis, raw MS spectra were processed by using Proteome Discoverer 2.4 software. SILAC 2plex (Arg10Lys6) method was selected to carry out the quantification analysis. The following search parameters were employed: full tryptic specificity was required, two missed cleavages were allowed; Carbamidomethylation was set as fixed modification, whereas Oxidation (M), Deamidation NQ), Acetylation (N-terminus) and GGE (K) were considered as variable modifications. Precursor ion mass tolerances were 10 ppm for all MS acquired and fragment ion mass tolerance was 0.02 Da for all MS2 spectra. The search results were automatically processed at the FDR 1% at both the protein and peptide. The unique peptide included in the protein group were used for quantification. Proteins with SILAC ratio greater than or equal 2 are considered candidate interacting proteins.

### Function analysis of the proteins identified in SPIDER

To analyze protein-protein interaction, the protein list was uploaded to STRING^3^ database (https://string-db.org). The list of protein interactions was imported into Cytoscape^4^ (Version 3.8.2) for network presentation. The plugin of MCODE was used for network clustering, in which node score cutoff was set by 0.2 and K-Core was set by 2. Gene Ontology analysis were performed by PANTHER online tools (http://www.pantherdb.org, February 2021). GO Terms enrichment analysis was performed using Fisher’s exact tests (False Discover Rate corrected *p* < 0.05, minimum two-fold enrichment) using the annotations of interactors. The statistical significance for overrepresentation test was used Fisher’s Exact test. Volcano plots of proteins identified in this study were generated using MetaboAnalyst 5.0 (https://www.metaboanalyst.ca/) (*p* < 0.05, Fold Change ≥1.5). Protein expression score of Omicron RBD interactors in human tissues was obtained from Human Protein Atlas database^5^ (www.proteinatlas.org). Protein expression of Omicron RBD interactors in cells from human tissues was analyzed using a single-cell mRNA sequencing dataset^6^ (http://bis.zju.edu.cn/HCL/).

### Structure analysis and sequence alignment

The nucleic acid and protein structures were obtained from PDB database (www.rcsb.org). Structural analysis was 0carried out by PyMOL (Version 2.5.0) at default setting. Sequence of Protein-g in different species was obtained from Uniprot (www.uniprot.org). Sequence alignment was performed by SnapGene (6.0).

### Bio-layer Interferometry

For measuring the binding kinetics, biotinylated protein was loaded at 25 ng/μL in kinetics buffer containing 1×PBS with 0.1% BSA and 0.02% Tween20 onto streptavidin biosensors (ForteBio, USA). Association of candidate protein was tested in kinetics buffer at gradient concentration for 3-5 min. Dissociation in kinetics buffer was measured for 3-5 min. BLI assays were carried out in 96-well black plates and analyzed on an OctetRed96 (Fortebio, USA) equipment. Mean K_on_, K_off_, K_D_ values were calculated by a 1:1 global fit model using the Data Analysis software (ForteBio, USA). The curves were processed using Prism software (GraphPad Prism 8.0.0).

### Immunofluorescence staining

HEK293T cells were seeded on poly-L-Lysine (Beyotime Biotechnology, China) coated coverslips and transfected with ACE2-Flag, PD-1-Flag, Flag-CCDC25 and Flag-tagged empty vector individually for 48 h. Cells were incubated with a wheat germ agglutinin (WGA) conjugate (Thermo, USA) with 5μg/mL for 15 min away from light at room temperature. After washed by PBS, cells were fixed with 4% paraformaldehyde for 15 min and blocked with 5% BSA in PBS for 30 min at room temperature. Cells were immunostained with an anti-Flag antibody (Sigma, USA) (1:100) overnight at 4 °C, followed by incubation with Alexa-488 conjugated anti-mouse antibody (Thermo, USA) (1:1000) for 1 h. The coverslips were mounted on microscope slides using Antifade Mounting Medium (Beyotime Biotechnology, China). Images were acquired with a confocal microscope (Leica SP8 STED).

### Flow cytometry

Cell suspensions were incubated in PBS with 2% FBS. 1 × 10^6 Cells were stained with PE anti-human CD11b antibodies (20 μL per test, BD Pharmingen, USA) and APC-Cy7 anti-human CD14 antibodies (5 μL per test, BD Pharmingen, USA) respectively for 30 min on ice. After stained with antibodies, cells were washed with PBS (2% FBS) twice by centrifuge at 600 rpm for 1 min. ~1 × 10^5 events were collected for each sample with a BD LSRFortessa system (BD Biosciences, USA) and FlowJo software 7.6.1 was used for data process.

### Structure modeling for the complex of human vimentin and Omicron RBD

The complex modeling process contained three steps, *i. e*., global docking, local docking, and the complex structure optimization.

A rough vimentin-RBD complex structure was obtained using a global rigid protein-protein docking program SDOCK^7^, which uses grid-based fast Fourier transform method for speed up the sampling of 3D translational docking space and uses stepwise force-field potentials for the scoring functions. The ACE2 binding structure of Omicron RBD (PDBID, 7WBP^8^, chain B, residues 333-527) and the dimer structure of human vimentin 2B fragment (PDBID, 1GK4^9^, residues 328-406 of chains A and B) were used as the input structures, and the default parameters were used for SDOCK docking. If a docking solution has at least 4 of the 15 RBD mutation sites of Omicron RBD compared to the Prototype were at the interface of RBD-vimentin, it was filtered out. The refinement of the global protein-protein docking model was achieved using RosettaDock^10^, which Monte Carlo simulations to search the rigid-body and side-chain conformation space of two interacting proteins to find a minimum free-energy complex structure. We assumed that the best complex model is not very far from the initial model, thus only 18 of the 200 models, which have a Ca-RMSD smaller than 4.0 Å to the initial model, were selected. Then, the Rosetta relax method^11^ was used to the model with the lowest total score. The Prototype RBD structure (PDB 6LZG^12^ chain B) was superposed to the vimentin - Omicron RBD complex structure to obtain the complex structure of vimentin - Prototype RBD complex structure, which was also optimized with Rosetta relax method. The residues 328-355 of vimentin rod chains A and B which were far from the binding interface were removed in the final models for molecular dynamics simulation and further analyses.

### The Molecular dynamic simulation of Protein-g-RBD complex for further structure relaxation and binding free energy estimation

The molecular dynamic simulations of the Protein-g - Omicron RBD complex or the Protein-g - Prototype RBD complex were conducted using AMBER18 MD simulation package with Amber ff19SB^13^ force field. Using the leap module, the complex was neutralized by adding sodium ions and was solved in a cubic TIP3P water box, and the distance between the box face and the solute was set to larger than 12 Å. Energy minimization was performed on the systems by steepest descent for 4000 steps, followed by conjugate gradient for the next 6000 steps. MD simulations were initiated by gradually heating each system in canonical ensemble (NVT) from 0 to 300 K in three steps for 380 picoseconds (ps). Langevin thermostate was used with coupling coefficient of 2 ps. Then, density equilibration was performed for 1 ns in isothermal-isobaric (NPT, 300 K, 1 bar) ensemble. Berendsen barostat with a pressure-coupling constant of 2 ps was used. The production MD simulations were carried out for 18 ns for each system in NPT (300 K, 1 bar) ensemble. During MD simulations, the particle mesh Ewald (PME) approach was used to treat long-range electrostatic interactions and van der Waals interactions were calculated with a cut-off of 12 Å. SHAKE algorithm was applied to constrain all the covalent bonds involving hydrogen atoms, and the time step was set to 2 femtoseconds and the structural coordinates were saved every 1 ps.

### Computational estimation of the binding free energies for omicron/ the Prototype RBD to Protein-g

The binding free energies of the two types of RBD to Protein-g were estimated based on trajectories of MD simulations by molecular mechanics/generalized Born surface area (MM/GBSA) method using the mmpbsa.py^14^ module in AMBER18 package^15^. We neglected the entropy term to compare the binding affinities of different RBD type to Protein-g and the single trajectory method^16^ was used. A total of 1500 snapshots were used for each system, which were from the production of MD simulation with an interval of 10 ps. The binding free energy was estimated by summing the gas phase binding free energy and the solvation free energy. The electrostatic solvation free energy was calculated using the generalized Born (GB) model. Omicron RBD + Protein-g: −49.7 ± 9.0 kcal/mol, the Prototype RBD + Protein-g: −15.7 ± 8.7 kcal/mol.

## Chemistry

**Scheme 1.**
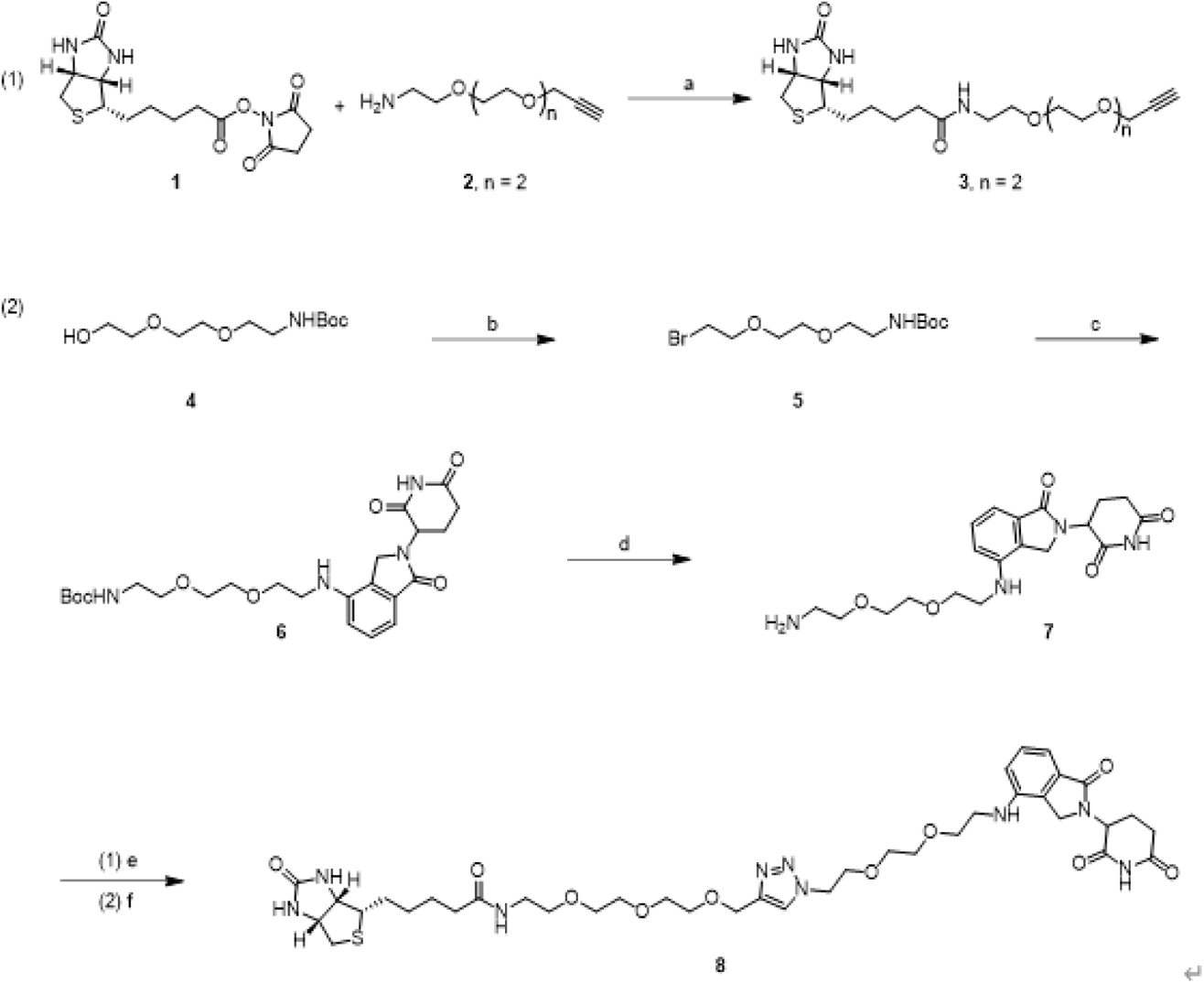
The synthesis of biotin labeled lenalidomide.

Reagent and conditions: (a) TEA, DMF, room temperature; (b) CBr_4_, PPh_3_, THF, room temperature; (c) lenalidomide, DIPEA, NMP, 110°C; (d) TFA, DCM, room temperature; (e) FSO_2_N_3_ (0.3M in MTBE), KHCO_3_ (2M in H_2_O), DMF, room temperature; (f) compound **3**, sodium ascorbate, CuSO_4_·5H_2_O, DMF, H_2_O, 50 °C.

## General

All reagents were commercially available and used without further purification. ^1^H nuclear magnetic resonance (NMR) spectra were recorded on an Agilent-400 instrument at 400 MHz. ^13^C NMR spectra were recorded on a Bruker AM-400 instrument at 101 MHz. Chemical shifts (δ) were expressed in parts per million (ppm) relative to residual solvent as an internal reference for ^1^H and ^13^C NMR. Coupling constants (*J*) were reported in hertz unit (Hz) and coupling patterns were described as singlet (s), doublet (d) and triplet (t). High resolution mass spectra (HRMS) were carried out on an Agilent Technologies 6230 LC-MS with (ESI-TOF) mode. Melting points (m.p.) were determined on a Büchi M-565 melting point apparatus. HPLC analysis of all final biological testing compounds was performed on a Waters ACQUITY UPLC H-Class system with a Waters ACQUITY QDa system operating in the electrospray ionization (ESI) mode with a flow rate of 0.6 mL/min and a gradient of eluting with H_2_O (with 0.1% trifluoroacetic acid) and CH_3_CN. The method used is as following: 7000 psi, flow rate = 0.6 ml/min, eluent: t = 0 min, 95% H_2_O; t = 0.5 min, 95% H_2_O; t = 3.5 min, 5% H_2_O; t = 4.5 min, 5% H_2_O; t = 5.0 min, 95% H_2_O, total acquisition time = 5.0 min. Purity of all final testing compounds was based on the integrated UV chromatogram at 220 nm.

### 5-((3aS,4S,6aR)-2-Oxohexahydro-1H-thieno[3,4-d]imidazol-4-yl)-N-(2-(2-(2-(prop-2-yn-1-yloxy)ethoxy)ethoxy)ethyl)pentanamide (3)

To a solution of biotin-NHS (**1**) (912 mg, 2.67 mmol) in DMF (10 mL) was added compound **2** (500 mg, 2.67 mmol) and TEA (689 μL, 5.34 mmol). The reaction mixture was stirred at room temperature overnight. The solvent was then removed *in vacuum* and the residue was purified by silica gel column chromatography (dichloromethane/methanol = 10/1) to afford compound **3** (943 mg, 85%) as a light yellow solid. ^1^H NMR (400 MHz, DMSO-*d_6_*): δ 7.84 (t, *J* = 5.6 Hz, 1H), 6.43 (s, 1H), 6.36 (s, 1H), 4.32-4.28 (m, 1H), 4.14-4.11 (m, 3H), 3.55-3.48 (m, 8H), 3.44 (t, *J* = 2.4 Hz, 1H), 3.39 (t, *J* = 6.0 Hz, 2H), 3.20-3.15 (m, 2H), 3.11-3.07 (m, 1H), 2.84-2.79 (m, 1H), 2.59-2.55 (m, 1H), 2.06 (t, *J* = 7.2 Hz, 2H), 1.65-1.22 (m, 6H) ppm; ^13^C NMR (101 MHz, DMSO-*d_6_*): δ 172.0, 162.6, 80.3, 77.0, 69.6, 69.5, 69.4, 69.1, 68.4, 60.9, 59.1, 57.4, 55.3, 39.7, 38.3, 35.0, 28.1, 27.9, 25.2 ppm.

### tert-Butyl (2-(2-(2-bromoethoxy)ethoxy)ethyl)carbamate (5)

To a solution of *tert*-butyl (2-(2-(2-hydroxyethoxy)ethoxy)ethyl)carbamate (1.0 g, 4.0 mmol) in THF (40 mL) was added CBr_4_ (2.7 g, 8.0 mmol) and PPh_3_ (2.1 g, 8 mmol). The reaction mixture was stirred at room temperature for 5 h and then filtered through celite. The filtrate was evaporated *in vacuum* and the residue was purified by silica gel column chromatography (petroleum ether/ethyl acetate = 5/1) to afford crude compound **5** (1.1 g) as a light yellow oil which was used in next step without further purification.

### tert-Butyl (2-(2-(2-((2-(2,6-dioxopiperidin-3-yl)-1-oxoisoindolin-4-yl)amino)ethoxy)ethoxy)ethyl) carbamate (6)

To a solution of lenalidomide (518 mg, 2 mmol) in NMP (8 mL) was added DIPEA (1 mL, 6 mmol) and compound **5** (936 mg, 3 mmol). Then the reaction mixture was heated to 110 °C for 12 h and monitored by LC-MS. The mixture was cooled to room temperature and solvent was evaporated *in vacuum*. The residue was purified by flash column (5% acetonitrile to 100% acetonitrile, 45 min) to afford compound **6** (307 mg, 31%) as a yellow solid. ^1^H NMR (400 MHz, DMSO-*d_6_*): δ 11.00 (s, 1H), 7.29 (t, *J* = 8.0 Hz, 1H), 6.94 (d, *J* = 7.6 Hz, 1H), 6.80 (d, *J* = 8.0 Hz, 1H), 6.77-6.72 (m, 1H), 5.59 (t, *J* = 5.2 Hz, 1H), 5.12 (d, *J* = 5.2 Hz, 0.5H), 5.09 (d, *J* = 4.8 Hz, 0.5H), 4.23 (d, *J* = 17.2 Hz, 1H), 4.12 (d, *J* = 17.2 Hz, 1H), 3.60-3.52 (m, 6H), 3.37 (t, *J* = 6.0 Hz, 2H), 3.33-3.22 (m, 2H), 3.08-3.03 (m, 2H), 2.97-2.88 (m, 1H), 2.63-2.59 (m, 1H), 2.36-2.25 (m, 1H), 2.05-2.01 (m, 1H), 1.36 (s, 9H) ppm; ^13^C NMR (101 MHz, DMSO-*d_6_*): δ 172.8, 171.1, 168.7, 155.5, 143.5, 132.0, 129.1, 126.4, 111.9,110.2, 77.5, 69.6, 69.4, 69.1, 68.8, 51.4, 45.6, 42.4, 31.2, 28.1, 22.7 ppm.

### 3-(4-((2-(2-(2-Aminoethoxy)ethoxy)ethyl)amino)-1-oxoisoindolin-2-yl)piperidine-2,6-dione (7)

To a solution of compound **6** (100 mg, 0.2 mmol) in dichloromethane (8 mL) was added TFA (2 mL). The reaction mixture was stirred at room temperature for 6 h and evaporated *in vacuum*. The residue was used for next step without further purification.

### N-(2-(2-(2-((1-(2-(2-(2-((2-(2,6-Dioxopiperidin-3-yl)-1-oxoisoindolin-4-yl)amino)ethoxy)ethoxy)ethyl)-1H-1,2,3-triazol-4-yl)methoxy)ethoxy)ethoxy)ethyl)-5-((3aS,4S,6aR)-2-oxohexahydro-1H-thieno[3,4-d]imidazol-4-yl)pentanamide (8)

To a solution of compound **7** (0.2 mmol) in DMF (2 mL) was added KHCO_3_ (2 M in H_2_O, 0.4 mL, 0.8 mmol) aqueous and FSO_2_N_3_ (0.3 M in MTBE, 0.8 mL, 0.24 mmol). After stirred at room temperature for 2 h, to the mixture was added sodium ascorbate (158 mg, 0.8 mmol), biotin-PEG3-alkyne **3** (82.6 mg, 0.2 mmol) and CuSO_4_·5H_2_O (10 mg, 0.04 mmol). The mixture was then heated to 50 °C. After stirred for 2 h, the mixture was cooled to room temperature and evaporated *in vacuum*. The residue was purified by flash column chromatography (0% acetonitrile, 100% H_2_O to 100% acetonitrile, 40 min) to afford compound **8** (87 mg, 52%) as a yellow solid. ^1^H NMR (400 MHz, DMSO-*d_6_*): δ 11.0 (s, 1H), 8.04 (s, 1H), 7.84 (t, *J* = 5.6 Hz, 1H), 7.29 (t, *J* = 7.6 Hz, 1H), 6.94 (d, *J* = 7.2 Hz, 1H), 6.79 (d, *J* = 8.0 Hz, 1H), 6.42 (s, 1H), 6.36 (s, 1H), 5.57 (t, *J* = 5.2 Hz, 1H), 5.12 (d, *J* = 4.8 Hz, 0.5H), 5.09 (d, *J* = 5.2 Hz, 0.5H), 4.51-4.46 (m, 4H), 4.31-4.28 (m, 1H), 4.22 (d, *J* = 17.2 Hz, 1H), 4.14-4.09 (m, 2H), 3.81 (t, *J* = 5.2 Hz, 2H), 3.56-3.48 (m, 14H), 3.39-3.36 (m, 2H), 3.32-3.27 (m, 2H), 3.19-3.15 (m, 2H), 3.10-3.06 (m, 1H), 2.97-2.88 (m, 1H), 2.83-2.79 (m, 1H), 2.63-2.55 (m, 2H), 2.35-2.24 (m, 1H), 2.07-2.01 (m, 3H), 1.64-1.25 (m, 6H) ppm; ^13^C NMR (101 MHz, DMSO-*d_6_*): δ 172.8, 172.0, 171.1, 168.7, 162.6, 143.7, 143.5, 132.0, 129.1, 126.4, 124.1, 111.9, 110.2, 69.6, 69.5, 69.4, 69.1, 68.9, 68.8, 68.6, 63.4, 60.9, 59.1, 55.3, 54.8, 51.4, 49.2, 45.6, 42.4, 39.7, 38.3, 35.0, 31.1, 28.1, 27.9, 25.2, 22.7 ppm. HRMS (ESI): [M+H]^+^ C_38_H_56_N_9_O_10_S calcd 830.3871, found 830.3865; m.p.: 40.5-43.6 °C; HPLC: purity 96.7%, retention time 2.10 min.

### N-(2-(2-(2-((1-((R)-4-Oxo-4-(3-(trifluoromethyl)-5,6-dihydro-[1,2,4]triazolo[4,3-a]pyrazin-7(8H)-yl)-1-(2,4,5-trifluorophenyl)butan-2-yl)-1H-1,2,3-triazol-4-yl)methoxy)ethoxy)ethoxy)ethyl)-5-((3aS,4S,6aR)-2-oxohexahydro-1H-thieno[3,4-d]imidazol-4-yl)pentanamide (12)

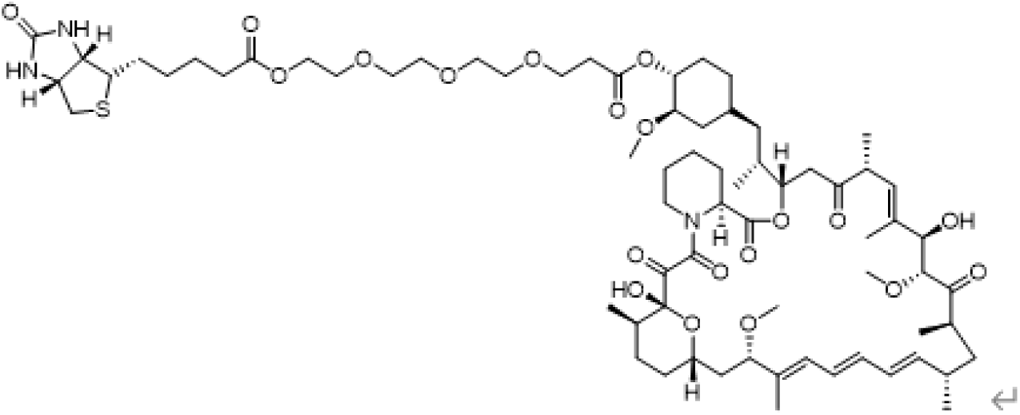

To a solution of rapamycin (Selleck) and biotin-PEG-acid in anhydrous CH_2_Cl_2_ were added DMAP, EDCI and Et_3_N at room temperature. After stirred for two days, the mixture was quenched with water, diluted with ethyl acetate, and washed with brine. The organic layer was filtered and concentrated. The resulting residue was purified by preparative thin-layer chromatography to provide the product.

### NMR spectra

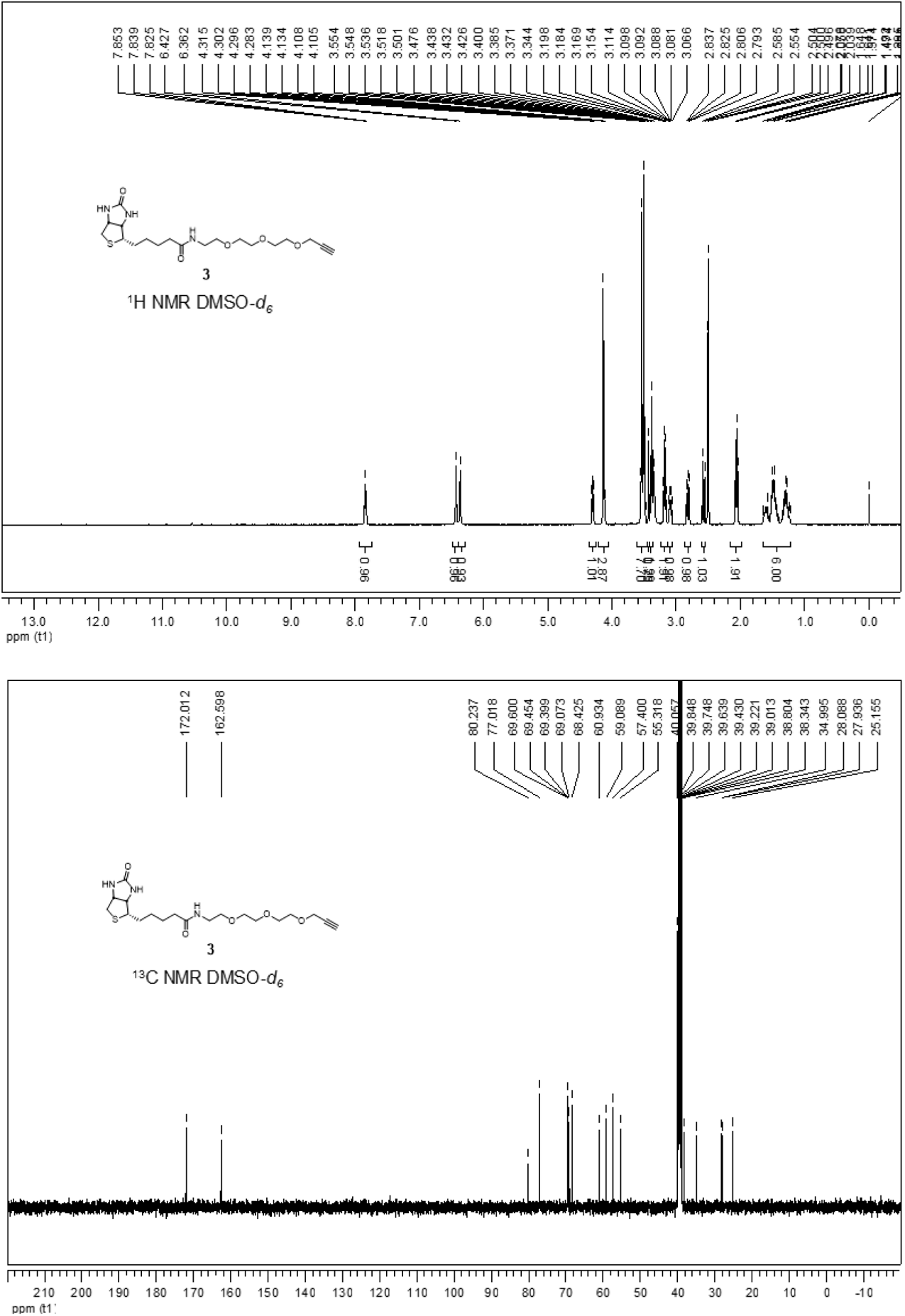
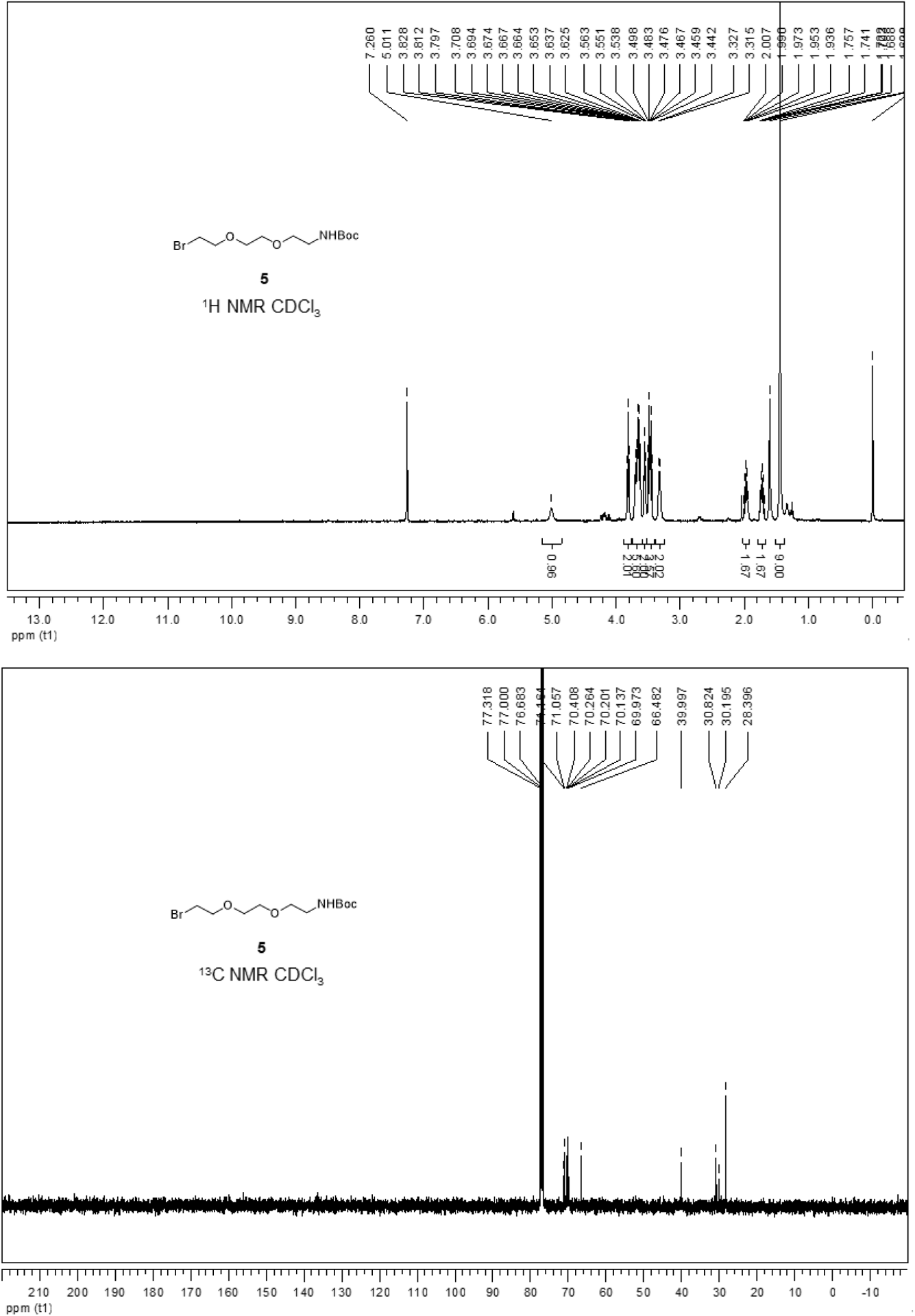
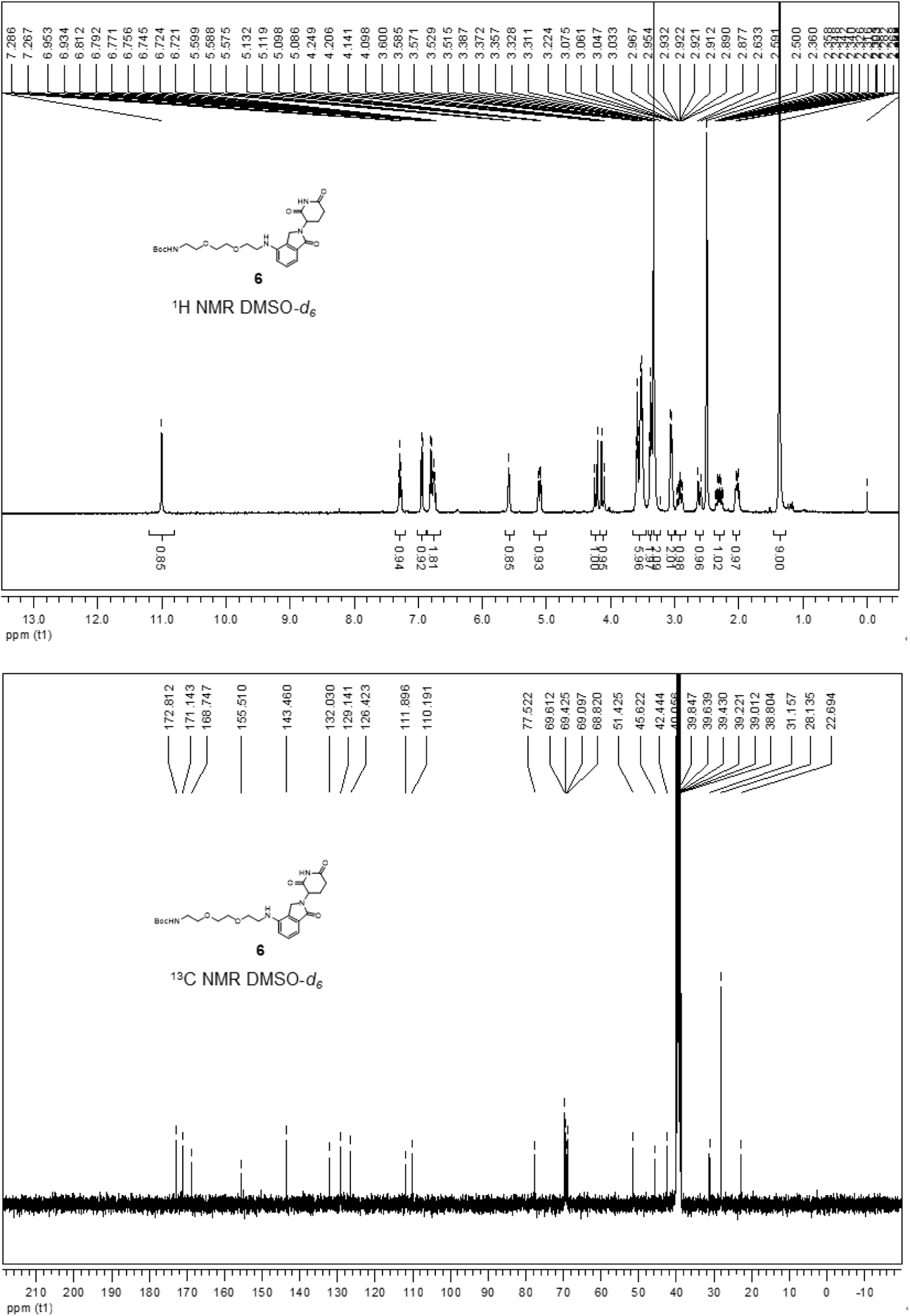
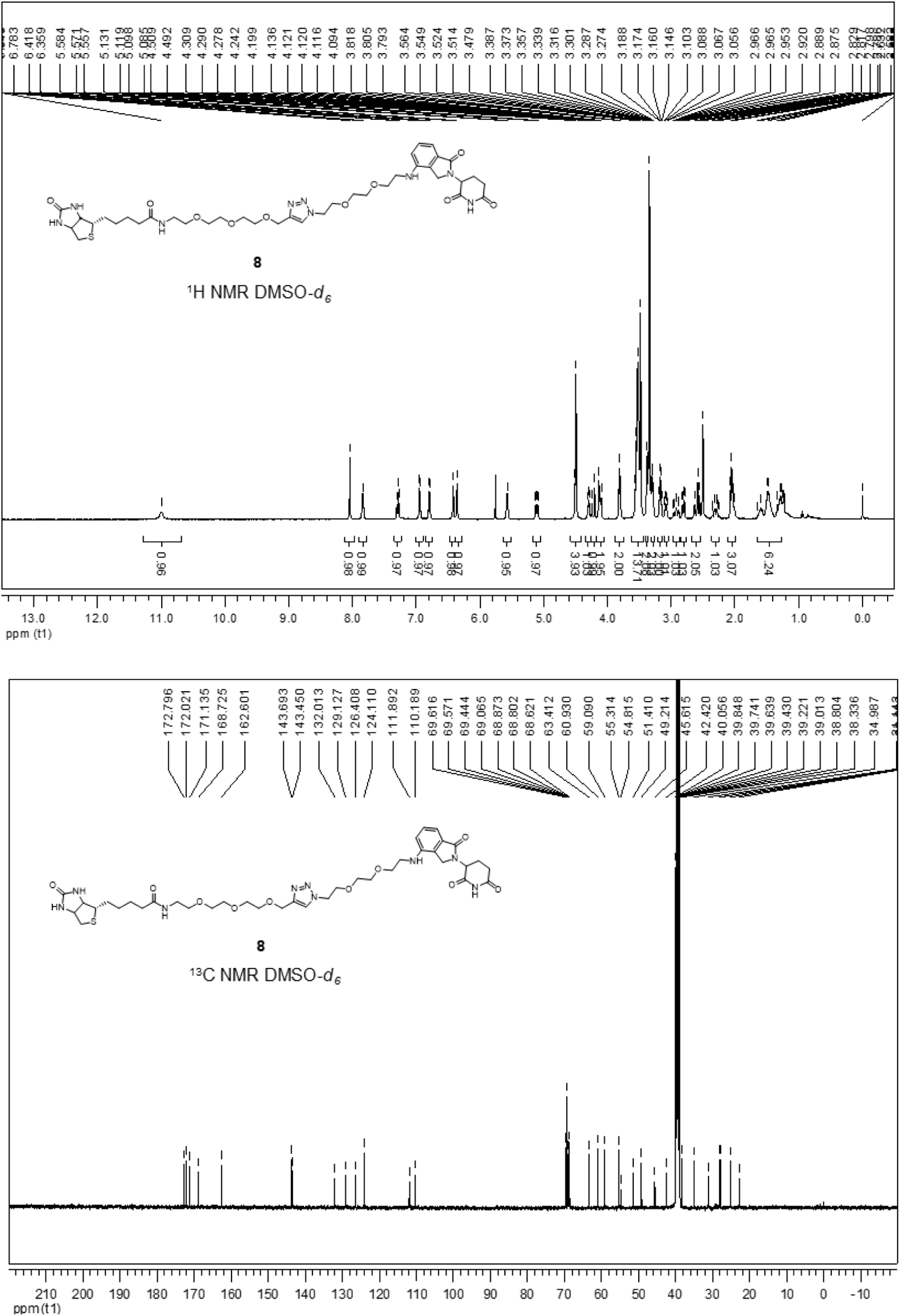

## Supplementary Tables

Supplementary Table 1. CobB interacting proteins identified by SPIDER.

Supplementary Table 2. CobB interacting proteins identified by SPIDER compared with two studies.

Supplementary Table 3. m6A binding proteins identified by SPIDER.

Supplementary Table 4. Lenalidomide interacting proteins identified by SPIDER.

Supplementary Table 5. Candidate receptors of SARS-CoV-2 identified by SPIDER.

Supplementary Table 6. Comparison of candidate receptors identified by SPIDER with BioGRID database.

Supplementary Table 7. Potential Omicron specific receptors.

Supplementary Table 8. Protein and mRNA expression score of candidate receptors enriched by Omicron RBD.

Supplementary Table 9. Detailed information of the plasmids and nucleic acid sequence prepared in this study.

## References

1. Ho, Y. et al. Systematic identification of protein complexes in Saccharomyces cerevisiae by mass spectrometry. Nature 415, 180–183,(2002).

2. Vikis, H. G. & Guan, K. L. Glutathione-S-transferase-fusion based assays for studying protein-protein interactions. Methods Mol Biol 261, 175–186,(2004).

3. Zhu, H. et al. Global analysis of protein activities using proteome chips. Science 293, 2101–2105,(2001).

4. Burckstummer, T. et al. An efficient tandem affinity purification procedure for interaction proteomics in mammalian cells. Nat Methods 3, 1013–1019,(2006).

5. Roux, K. J. et al. A promiscuous biotin ligase fusion protein identifies proximal and interacting proteins in mammalian cells. J Cell Biol 196, 801–810,(2012).

6. Rhee, H. W. et al. Proteomic mapping of mitochondria in living cells via spatially restricted enzymatic tagging. Science 339, 1328–1331,(2013).

7. Liu, Q. et al. A proximity-tagging system to identify membrane protein-protein interactions. Nat Methods 15, 715–722,(2018).

8. Qin, W. et al. Deciphering molecular interactions by proximity labeling. Nat Methods 18, 133–143,(2021).

9. Luscombe, N. M. et al. An overview of the structures of protein-DNA complexes. Genome Biol 1, REVIEWS001,(2000).

10. Van Nostrand, E. L. et al. A large-scale binding and functional map of human RNA-binding proteins. Nature 583, 711–719,(2020).

11. Gerstberger, S. et al. A census of human RNA-binding proteins. Nat Rev Genet 15, 829–845,(2014).

12. Castello, A. et al. Insights into RNA biology from an atlas of mammalian mRNA-binding proteins. Cell 149, 1393–1406,(2012).

13. Liu, L. et al. Insight into novel RNA-binding activities via large-scale analysis of lncRNA-bound proteome and IDH1-bound transcriptome. Nucleic Acids Res 47, 2244–2262,(2019).

14. Liu, X. et al. In Situ Capture of Chromatin Interactions by Biotinylated dCas9. Cell 170, 1028–1043 e1019,(2017).

15. Myers, S. A. et al. Discovery of proteins associated with a predefined genomic locus via dCas9-APEX-mediated proximity labeling. Nat Methods 15, 437–439,(2018).

16. Yi, W. et al. CRISPR-assisted detection of RNA-protein interactions in living cells. Nat Methods 17, 685–688,(2020).

17. Zhang, Z. et al. Capturing RNA-protein interaction via CRUIS. Nucleic Acids Res 48, e52,(2020).

18. Manghwar, H. et al. CRISPR/Cas Systems in Genome Editing: Methodologies and Tools for sgRNA Design, Off-Target Evaluation, and Strategies to Mitigate Off-Target Effects. Adv Sci (Weinh) 7, 1902312,(2020).

19. Bedard, P. L. et al. Small molecules, big impact: 20 years of targeted therapy in oncology. Lancet 395, 1078–1088,(2020).

20. Deshaies, R. J. Multispecific drugs herald a new era of biopharmaceutical innovation. Nature 580, 329–338,(2020).

21. Wishart, D. S. Metabolomics for Investigating Physiological and Pathophysiological Processes. Physiol Rev 99, 1819–1875,(2019).

22. Svatos, A. Mass spectrometric imaging of small molecules. Trends Biotechnol 28, 425–434,(2010).

23. Kawatani, M. & Osada, H. Affinity-based target identification for bioactive small molecules. Medchemcomm 5, 277–287,(2014).

24. Tsushima, M. et al. Selective purification and chemical labeling of a target protein on ruthenium photocatalyst-functionalized affinity beads. Chem Commun (Camb) 53, 4838–4841,(2017).

25. Park, J. et al. From noncovalent to covalent bonds: a paradigm shift in target protein identification. Mol Biosyst 9, 544–550,(2013).

26. Backus, K. M. et al. Proteome-wide covalent ligand discovery in native biological systems. Nature 534, 570–574,(2016).

27. Kulkarni, P. M. et al. Novel Electrophilic and Photoaffinity Covalent Probes for Mapping the Cannabinoid 1 Receptor Allosteric Site(s). J Med Chem 59, 44–60,(2016).

28. Mackinnon, A. L. & Taunton, J. Target Identification by Diazirine Photo-Cross-linking and Click Chemistry. Curr Protoc Chem Biol 1, 55–73,(2009).

29. Pearce, M. J. et al. Ubiquitin-like protein involved in the proteasome pathway of Mycobacterium tuberculosis. Science 322, 1104–1107,(2008).

30. Liao, S. et al. Pup, a prokaryotic ubiquitin-like protein, is an intrinsically disordered protein. Biochem J 422, 207–215,(2009).

31. Kim, J. et al. Use of in vivo biotinylation to study protein-protein and protein-DNA interactions in mouse embryonic stem cells. Nat Protoc 4, 506–517,(2009).

32. Wang, H. & Matsumura, P. Characterization of the CheAS/CheZ complex: a specific interaction resulting in enhanced dephosphorylating activity on CheY-phosphate. Mol Microbiol 19, 695–703,(1996).

33. Cantwell, B. J. & Manson, M. D. Protein domains and residues involved in the CheZ/CheAS interaction. J Bacteriol 191, 5838–5841,(2009).

34. Shimada, T. et al. The Escherichia coli RutR transcription factor binds at targets within genes as well as intergenic regions. Nucleic Acids Res 36, 3950–3955,(2008).

35. Hou, L. et al. Concurrent binding to DNA and RNA facilitates the pluripotency reprogramming activity of Sox2. Nucleic Acids Res 48, 3869–3887,(2020).

36. Dinesh, D. C. et al. Structural basis of RNA recognition by the SARS-CoV-2 nucleocapsid phosphoprotein. PLoS Pathog 16, e1009100,(2020).

37. Itoh, S. & Navia, M. A. Structure comparison of native and mutant human recombinant FKBP12 complexes with the immunosuppressant drug FK506 (tacrolimus). Protein Sci 4, 2261–2268,(1995).

38. Starai, V. J. et al. Sir2-dependent activation of acetyl-CoA synthetase by deacetylation of active lysine. Science 298, 2390–2392,(2002).

39. Weinert, B. T. et al. Acetyl-phosphate is a critical determinant of lysine acetylation in E. coli. Mol Cell 51, 265–272,(2013).

40. Castano-Cerezo, S. et al. Protein acetylation affects acetate metabolism, motility and acid stress response in Escherichia coli. Mol Syst Biol 10, 762,(2014).

41. Cheng, Z. F. & Deutscher, M. P. Purification and characterization of the Escherichia coli exoribonuclease RNase R. Comparison with RNase II. J Biol Chem 277, 21624–21629,(2002).

42. Liang, W. et al. Acetylation regulates the stability of a bacterial protein: growth stage-dependent modification of RNase R. Mol Cell 44, 160–166,(2011).

43. Huang, H. et al. Recognition of RNA N(6)-methyladenosine by IGF2BP proteins enhances mRNA stability and translation. Nat Cell Biol 20, 285–295,(2018).

44. Wang, X. et al. N6-methyladenosine-dependent regulation of messenger RNA stability. Nature 505, 117–120,(2014).

45. Xiao, W. et al. Nuclear m(6)A Reader YTHDC1 Regulates mRNA Splicing. Mol Cell 61, 507–519,(2016).

46. Fischer, E. S. et al. Structure of the DDB1-CRBN E3 ubiquitin ligase in complex with thalidomide. Nature 512, 49–53,(2014).

47. Chen, N. et al. Clinical Pharmacokinetics and Pharmacodynamics of Lenalidomide. Clin Pharmacokinet 56, 139–152,(2017).

48. Cao, Y. et al. Omicron escapes the majority of existing SARS-CoV-2 neutralizing antibodies. Nature 602, 657–663,(2022).

49. Han, P. et al. Receptor binding and complex structures of human ACE2 to spike RBD from omicron and delta SARS-CoV-2. Cell 185, 630–640 e610,(2022).

50. Xue, J. B. et al. Landscape of the RBD-specific IgG, IgM, and IgA responses triggered by the inactivated virus vaccine against the Omicron variant. Cell Discov 8, 15,(2022).

51. Meng, B. et al. Altered TMPRSS2 usage by SARS-CoV-2 Omicron impacts tropism and fusogenicity. Nature 10.1038/s41586-022-04474-x,(2022).

52. Cui, Z. et al. Structural and functional characterizations of infectivity and immune evasion of SARS-CoV-2 Omicron. Cell 10.1016/j.cell.2022.01.019,(2022).

53. Wang, S. et al. AXL is a candidate receptor for SARS-CoV-2 that promotes infection of pulmonary and bronchial epithelial cells. Cell Res 31, 126–140,(2021).

54. Uhlen, M. et al. Proteomics. Tissue-based map of the human proteome. Science 347, 1260419,(2015).

55. Han, X. et al. Construction of a human cell landscape at single-cell level. Nature 581, 303–309,(2020).

56. Shuai, H. et al. Attenuated replication and pathogenicity of SARS-CoV-2 B.1.1.529 Omicron. Nature 10.1038/s41586-022-04442-5,(2022).

57. Barandun, J. et al. Crystal structure of the complex between prokaryotic ubiquitin-like protein and its ligase PafA. J Am Chem Soc 135, 6794–6797,(2013).

## Reference

1. Ma, J. et al. iProX: an integrated proteome resource. Nucleic Acids Res 47, D1211–D1217,(2019).

2. Cox, J. & Mann, M. MaxQuant enables high peptide identification rates, individualized p.p.b.-range mass accuracies and proteome-wide protein quantification. Nature Biotechnology 26, 1367–1372,(2008).

3. Szklarczyk, D. et al. STRING v11: protein-protein association networks with increased coverage, supporting functional discovery in genome-wide experimental datasets. Nucleic Acids Res 47, D607–D613,(2019).

4. Shannon, P. et al. Cytoscape: a software environment for integrated models of biomolecular interaction networks. Genome Res 13, 2498–2504,(2003).

5. Uhlen, M. et al. Proteomics. Tissue-based map of the human proteome. Science 347, 1260419,(2015).

6. Han, X. et al. Construction of a human cell landscape at single-cell level. Nature 581, 303–309,(2020).

7. Zhang, C. & Lai, L. SDOCK: a global protein-protein docking program using stepwise force-field potentials. J Comput Chem 32, 2598–2612,(2011).

8. Han, P. Receptor binding and complex structures of human ACE2 to spike RBD from Omicron and Delta SARS-CoV-2. Cell, https://doi.org/10.1016/j.cell.2022.1001.1001,(2022).

9. Strelkov, S. V. et al. Conserved segments 1A and 2B of the intermediate filament dimer: their atomic structures and role in filament assembly. EMBO J 21, 1255–1266,(2002).

10. Lyskov, S. & Gray, J. J. The RosettaDock server for local protein-protein docking. Nucleic Acids Res 36, W233–238,(2008).

11. Khatib, F. et al. Algorithm discovery by protein folding game players. Proc Natl Acad Sci U S A 108, 18949–18953,(2011).

12. Wang, Q. et al. Structural and Functional Basis of SARS-CoV-2 Entry by Using Human ACE2. Cell 181, 894–904 e899,(2020).

13. Tian, C. et al. ff19SB: Amino-Acid-Specific Protein Backbone Parameters Trained against Quantum Mechanics Energy Surfaces in Solution. J Chem Theory Comput 16, 528–552,(2020).

14. Miller, B. R., 3rd et al. MMPBSA.py: An Efficient Program for End-State Free Energy Calculations. J Chem Theory Comput 8, 3314–3321,(2012).

15. Case, D. A. et al. The Amber biomolecular simulation programs. J Comput Chem 26, 1668–1688,(2005).

16. Gohlke, H. & Case, D. A. Converging free energy estimates: MM-PB(GB)SA studies on the protein-protein complex Ras-Raf. J Comput Chem 25, 238–250,(2004).

