## Supplemental Figure for "Specific Pupylation as IDEntity Reporter (SPIDER) for the identification of Protein-Biomolecule interactions"

### Supplementary Information

#### Identification of protein-biomolecule interactions by a substrate-based proximity labeling system

**Jiang *et al.***

**Table of contents**

**Extended Data Fig.1** Design and validation of the SPIDER proximity-tagging system

**Extended Data Fig.2** SPIDER to validate protein-biomolecule interaction

**Extended Data Fig.3** SPIDER to capture protein-protein interactions

**Extended Data Fig.4** SPIDER to capture protein-nucleic acid interactions

**Extended Data Fig.5** SPIDER to capture protein-small molecule interactions

**Extended Data Fig.6** SPIDER for identification of SARS-CoV-2 receptor

**Extended Data Fig.7** The SARS-CoV-2 Spike interacts with Protein-g

**Extended Data Fig.8** Structural analysis of RBD and Protein-g interaction

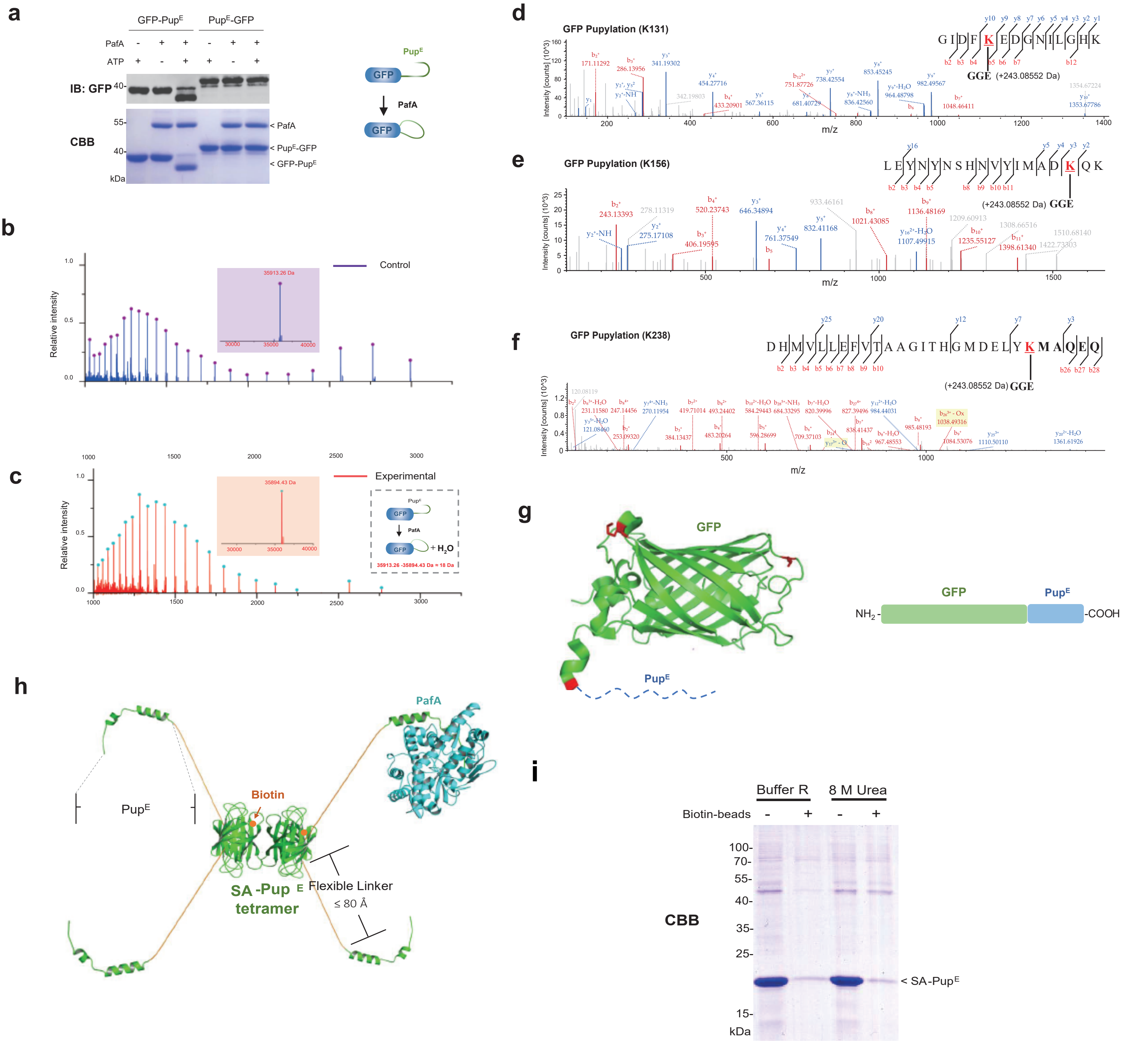


**Extended Data Fig.1 Design and validation of the SPIDER proximity-tagging system**

**Extended Data Fig.1 Design and validation of the SPIDER proximity-tagging system, related to Fig.1**

**a,** Self-pupylation of GFP-$Pup^E$ but not $Pup^E$-GFP. Reactions without PafA or ATP were included as controls. **b,c,** Characterization of GFP-Pup after Pupylation assays by mass spectrometry. **d-f,** Pupylation sites on GFP after SPIDER assay were identified by LC-MS/MS analysis, by searching for the additional mass of ~243.09 Da, K131(**d**), K156(**e**) and K238(**f**). **g,** Pupylation sites mapped on the crystal structure of GFP (PDB: 1B9C). The pupylated lysines were highlighted in red on the sequence. **h**, The model structure of SA-$Pup^E$~PafA. The 3D structure of SA was from M. T. Jacobsen *et al.*,[1]. The 3D structure of *Mtb* PafA is a homology modeling structure of *Corynebacterium glutamicum* PafA[2]. **i**, SDS-PAGE and Coomassie staining of SA-$Pup^E$ after depletion by biotin-agarose. After incubation with SA-$Pup^E$, the biotin-beads were washed by the SPIDER reaction buffer (Buffer R) or wash buffer with 8 M Urea. The beads were removed by centrifugation, the supernatants were visualized by SDS-PAGE and Coomassie staining.

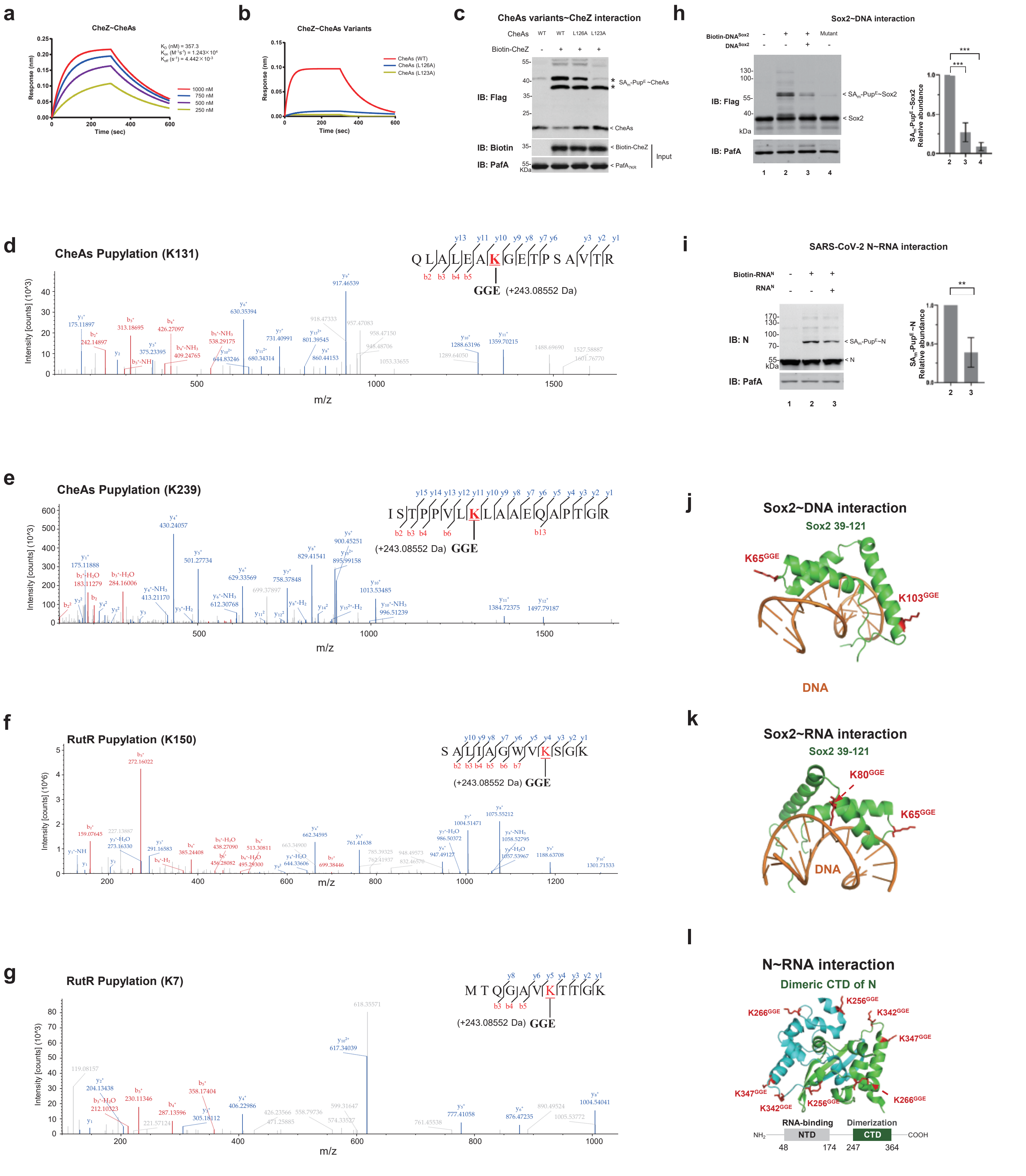


### Extended Data Fig.2 SPIDER to validate protein-based interaction

**Extended Data Fig.2 SPIDER to validate protein-biomolecule interaction, related to Fig.1**

**a,** Binding affinity of CheZ and CheAs (WT) was measured by BLI. **b,** Binding affinity of the two mutants of CheAs, *i. e.*, L126A, L123A. **c,** SPIDER assays with Flag-tagged CheAs mutants (L126A and L123A) and biotin-CheZ. **d,e,** Pupylation sites on WT CheAs after SPIDER assay. These sites were identified by LC-MS/MS analysis, by searching for an additional mass of ~243.09 Da. K131(**d**) and K239(**e**). **f,g,** Pupylation sites on WT RutR after SPIDER assay were identified by LC-MS/MS analysis. K150(**f**) and K7(**g**). **h,i,** SPIDER assays with Flag-tagged Sox2 and specific biotin-DNA (**h**). SPIDER assays with Flag-tagged SARS-CoV-2 N and specific biotin-RNA(**i**). Quantitation of Relative abundance of the major band of SA-Pup$^E$ ~ Prey based on western-blotting is shown at the right panel. Data are representative of one experiment with at least three independent biological replicates (mean and S.E.M. of n = 3), ** $p$ < 0.01 and *** $p$ < 0.001 (two-tailed unpaired t-test). Nucleic acid sequences can be found in Supplementary Table 6. **j-l,** Pupylation sites were mapped on the crystal structures of proteins, *i. e.*, Sox2 (PDB: 1O4X) (**j, k**), SARS-CoV-2 N (PDB: 7C22) (**i**). The pupylated lysines are highlighted in red.

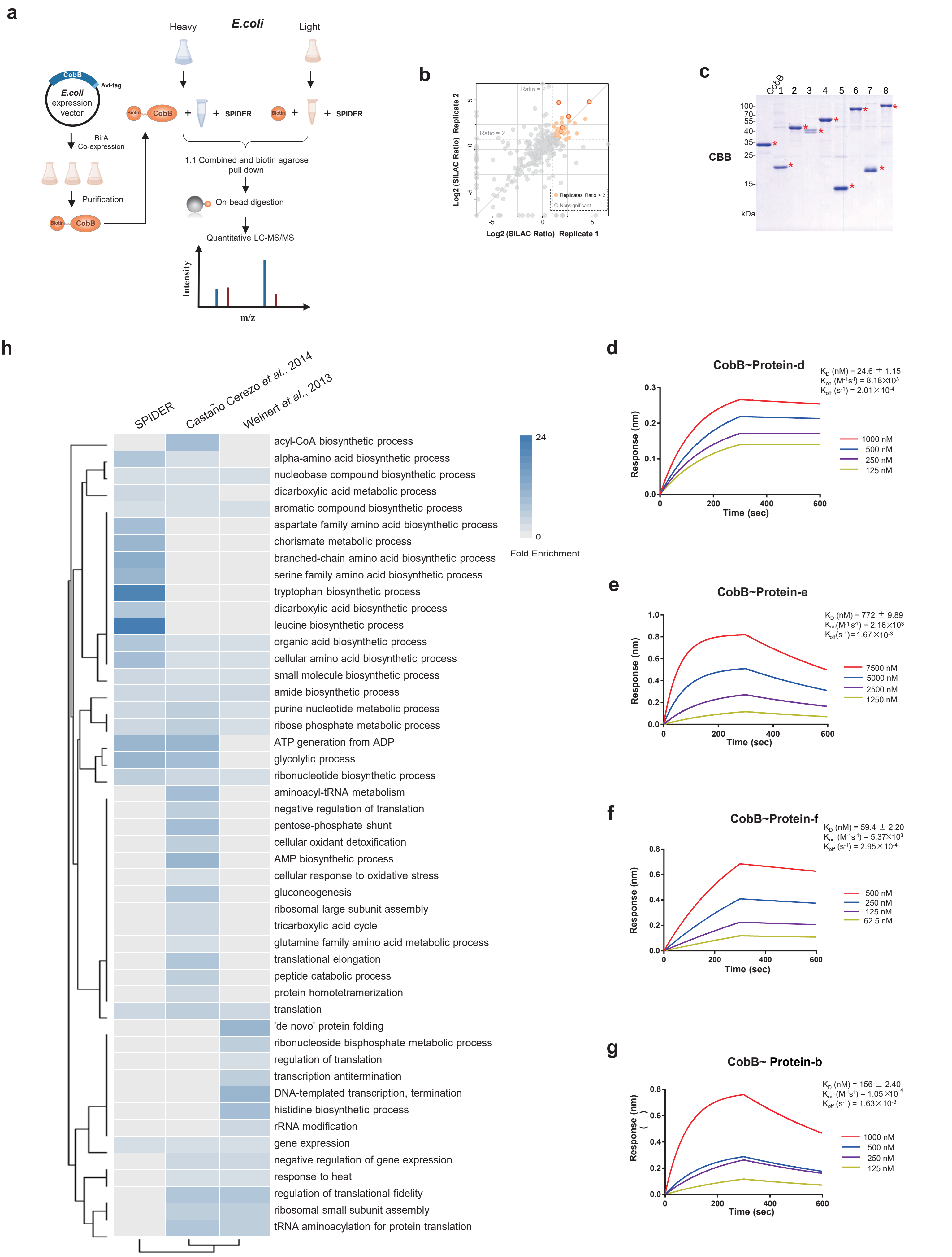


**Extended Data Fig.3 SPIDER to capture protein-protein interactions**

**Extended Data Fig.3 SPIDER to capture protein-protein interactions, related to Fig.2**

**a,** The workflow of SPIDER assays for capturing CobB interacting proteins. Biotinylated CobB was incubated with E. coli total lysate labeled with heavy stable isotope. As a control, free biotin was incubated with E. coli total lysate labeled with light stable isotope. **b,** Scatter plots show comparison of SILAC ratio of two replicate experiments of SPIDER~Biotin-CobB. The axes represent the log2 fold change (SILAC ratio) of protein quantification intensity (SPIDER~Biotin-CobB / SPIDER~Biotin). Significantly enriched proteins are shown as orange dots. **c,** Representative CobB interacting proteins identified by SPIDER. **d to g,** Binding affinity of CobB and four E. coli proteins, i. e., Protein-d**(d)**, Protein-e**(e)**, Protein-f**(f)** and Protein-b**(g)**, were measured using BLI. **h,** The CobB interacting proteins identified by SPIDER were compared with two other studies[3,4]. The top GO term of each group was selected for visualization (Fisher's exact test, one tailed). The number of proteins for each individual term is also shown.

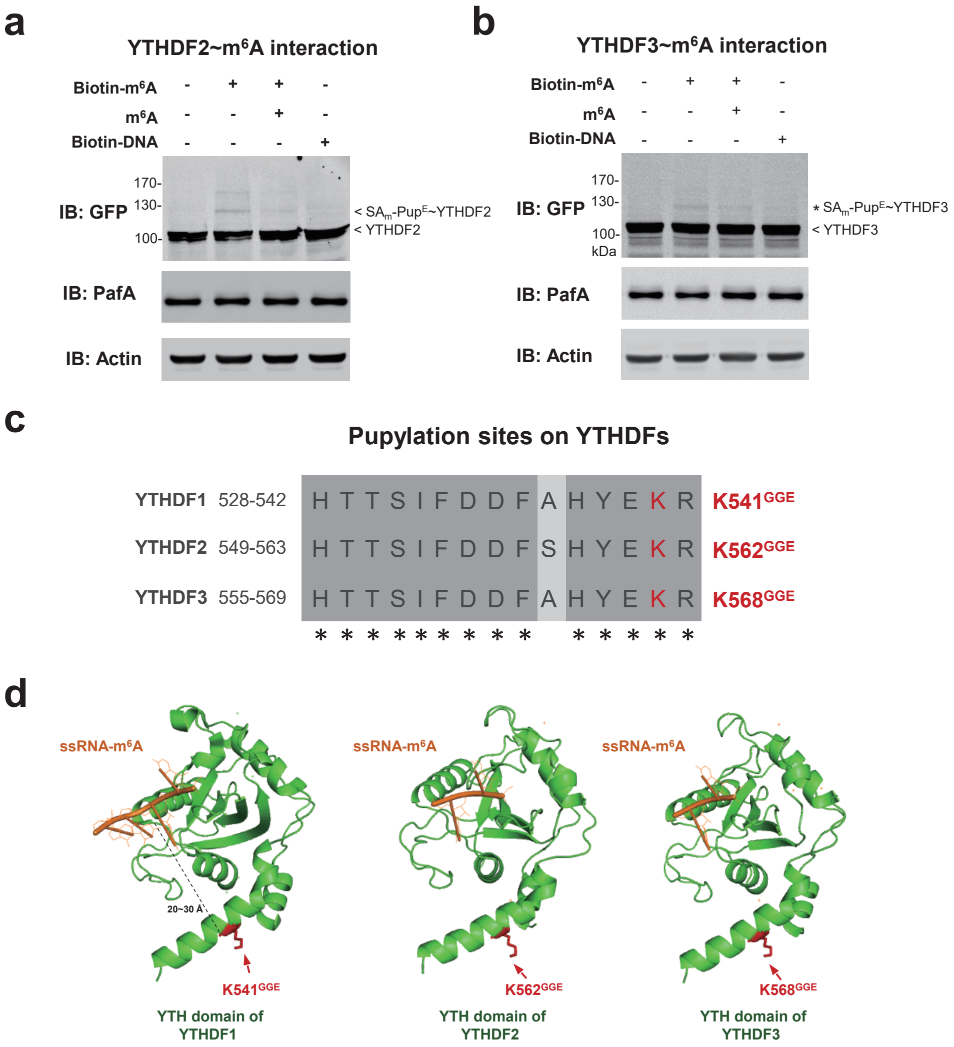


**Extended Data Fig.4 SPIDER to capture protein- nucleic acid interactions**

**Extended Data Fig.4 SPIDER to capture protein-nucleic acid interactions, related to Fig.3**

**a,b,** SPIDER assay with the cell lysate of YTHDF2 or YTHDF3- overexpressed HEK293T cells and biotin-m6A. RNA sequences can be found in Table S1. **c,** Alignment of the pupylated sites on 3 YTHDFs. The modified lysines are highlighted in red. **d,** Pupylated sites were mapped onto the crystal structure of YTHDFs YTH domain in complex with RNA. YTHDF1: PDB: 4RCJ. YTHDF2 or YTHDF3: The 3D structure is a homology modeling structure of YTHDF1. The modified lysines are highlighted in red.

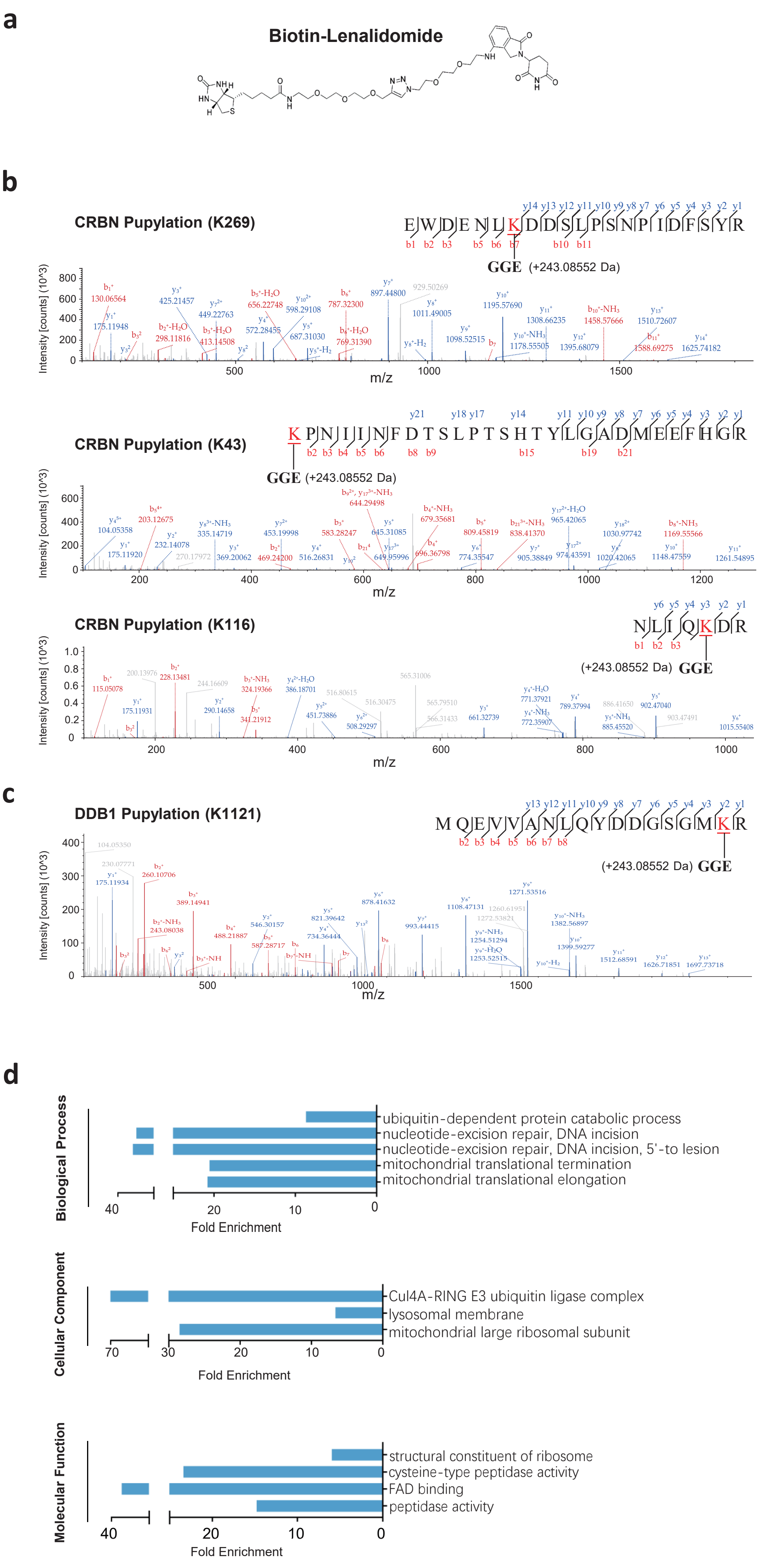


**Extended Data Fig.5 SPIDER to capture protein- small molecule interactions**

**Extended Data Fig.5 SPIDER to capture protein-small molecule interactions, related to Fig.4**

**a,** Chemical structure of biotin-Lenalidomide. **b,c,** Pupylation sites on CRBN or DDB1 were identified by LC-MS/MS analysis. K269, K43, K116**(b)**. K1121 **(c)**. **d,** Gene ontology analysis of the interacting proteins of Lenalidomide identified by SPDIER. The significantly enriched GO terms of each group were selected for visualization (Fisher's exact test, one tailed). The fold enrichment was given.

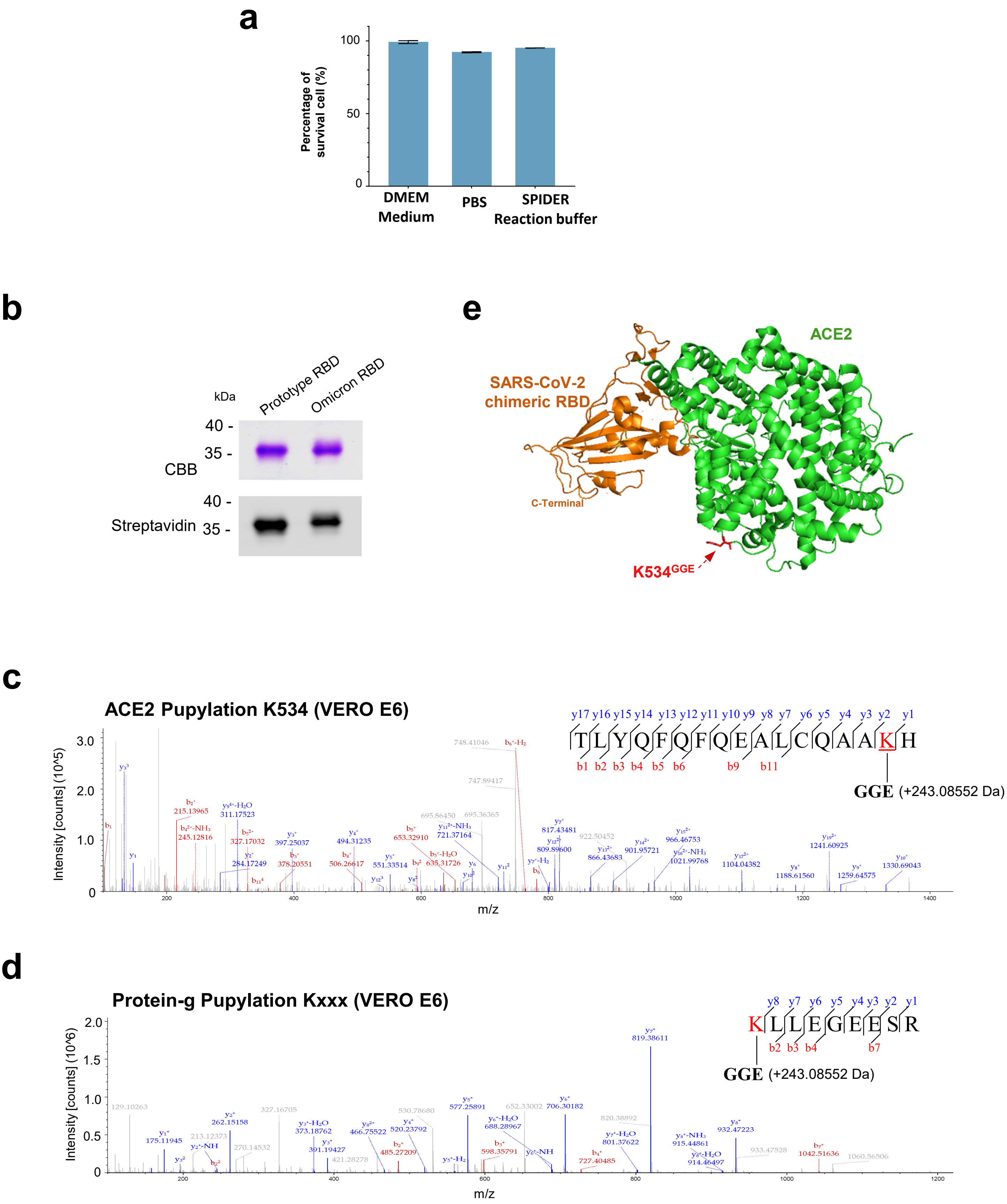


**Extended Data Fig.6 SPIDER for identification of SARS-CoV-2 receptor on cell surface**

**Extended Data Fig.6 SPIDER for identification of SARS-CoV-2 receptor, related to Fig. 5**

**a,** Percentage of survived HEK293T cell after 2 h incubation in different buffers. **b,** Biotinylated Prototype and Omicron RBDs were analyzed by SDS-PAGE, followed by Coomassie brilliant blue (CBB) staining and western blotting with Streptavidin. **c,** Pupylation site on ACE2 after SPIDER assay was identified by LC-MS/MS analysis, by searching for an additional mass of ~243.09 Da. **d,** Pupylation site (Kxxx) on Protein-g after SPIDER assay was identified by LC-MS/MS analysis, by searching for an additional mass of ~243.09 Da. **e,** Pupylation sites mapped on crystal structure of RBD-ACE2 complex (PDB: 6VW1). The pupylated lysine is highlighted in red.

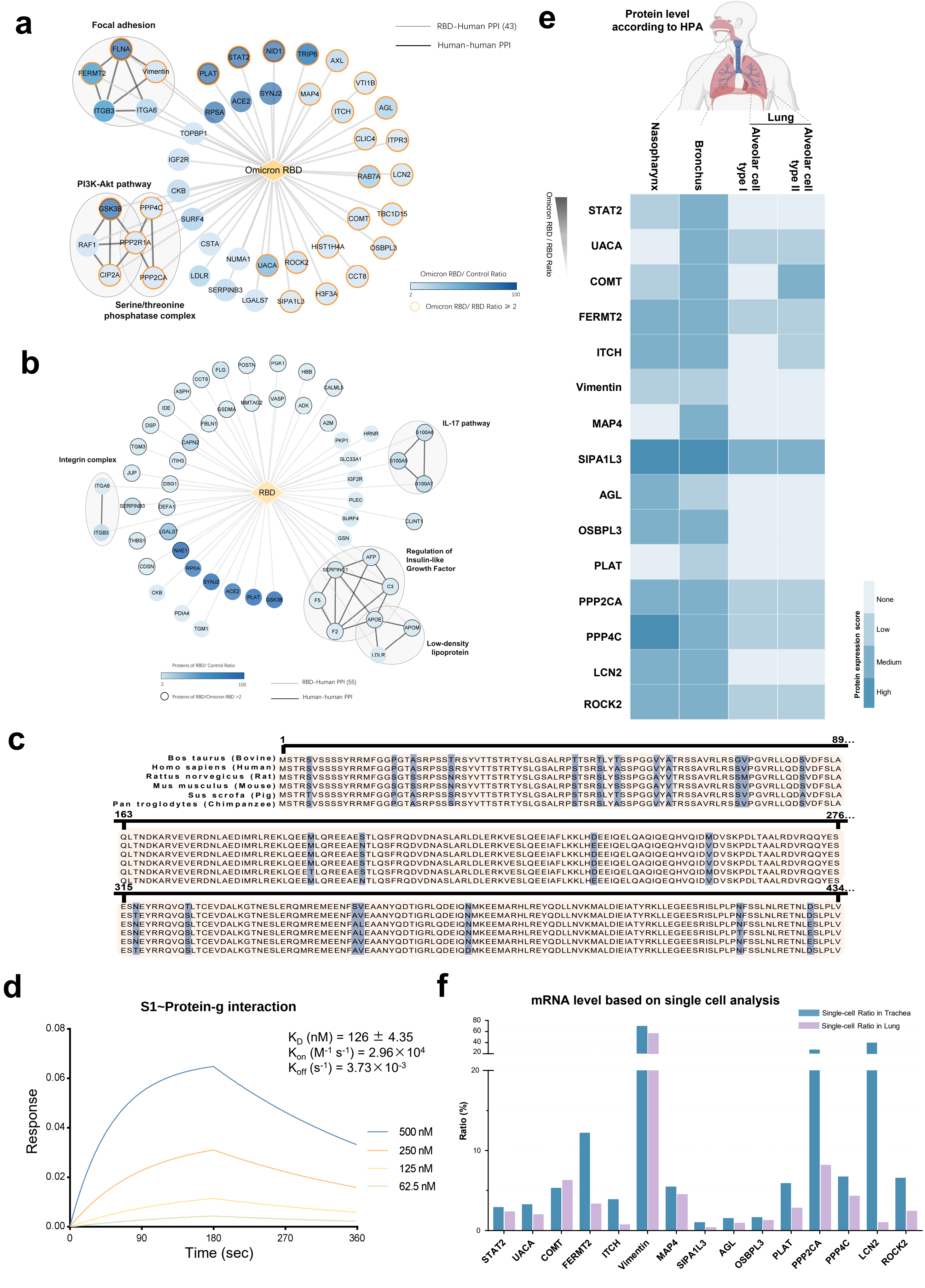


**Extended Data Fig.7 The SARS-CoV-2 Spike interacts with Protein-g**

**Extended Data Fig.7 The SARS-CoV-2 Spike interacts with Protein-g, related to Fig.5**

**a,** The network of the Omicron RBD interacting proteins. The interacting proteins identified by SPIDER were used as input. The interaction between human proteins were obtained from STRING with a confidence score > 0.7. **b,** The network of the Prototype RBD interacting proteins. The 55 interacting proteins identified by SPIDER as input. The interaction between human proteins were obtained from STRING with a confidence score > 0.7. **c,** Sequence alignment of Protein-g among species. Substituted residues are labeled with blue. **d,** Binding affinity of Prototype S1~Protein-g was measured using BLI. **e,** Heatmap of the protein expression score of proteins enriched by Omicron RBD. Protein expression score is obtained from Human Protein Atlas database (www.proteinatlas.org). **f,** The single-cell ratio of mRNA expression in trachea and lung. mRNA expression score is obtained from a single-cell mRNA sequencing dataset (http://bis.zju.edu.cn/HCL/).

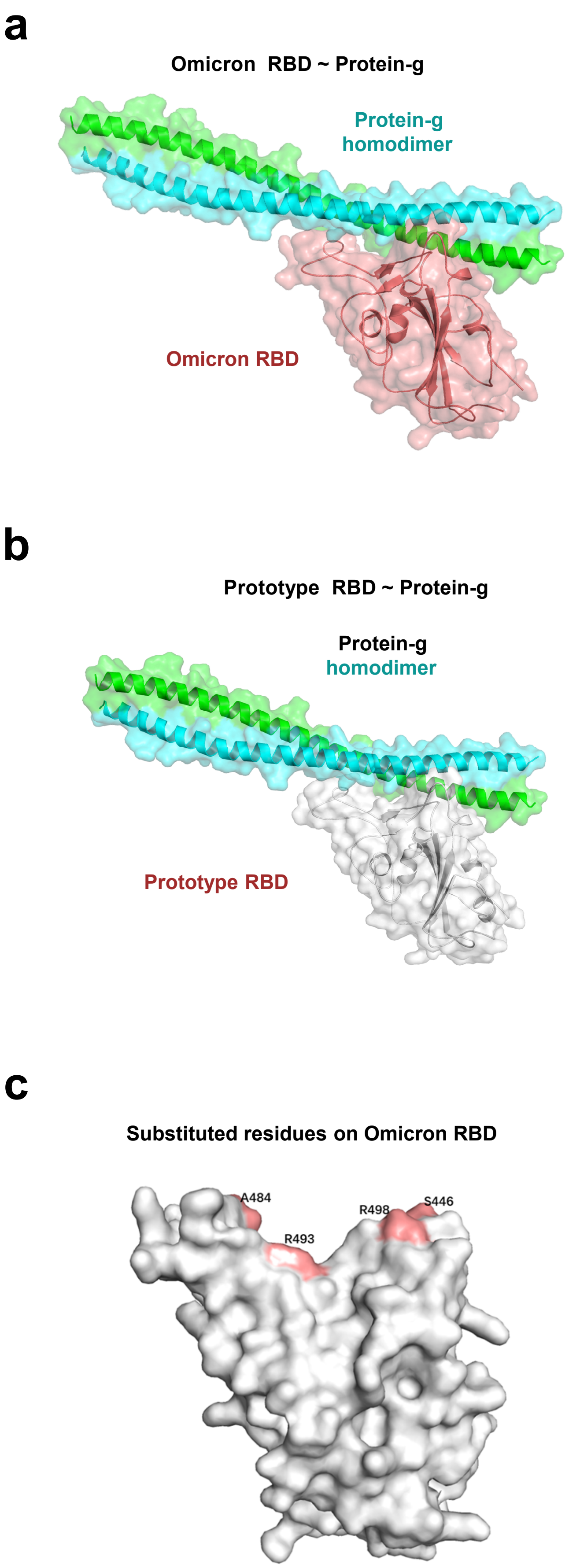


**Extended Data Fig.8 Structural analysis of RBD and Protein-g interaction**

**Extended Data Fig.8 Structural analysis of RBD and Protein-g interaction, related to Fig.5**

**a,b,** The predicted complex structures of Protein-g-Omicron RBD (**a**) and Protein-g-Prototype RBD (**b**). **c**,The binding surface of the Omicron RBD with Protein-g. Four different residues on Protein-g-binding interface of the Omicron RBD as compared to that of the Prototype RBD were labeled.

#### Reference


1 Jacobsen, M. T. *et al.* Amine Landscaping to Maximize Protein-Dye Fluorescence and Ultrastable Protein-Ligand Interaction. *Cell Chem Biol* **24**, 1040-1047 e1044,(2017).

2 Barandun, J. *et al.* Crystal Structure of the Complex between Prokaryotic Ubiquitin-like Protein and Its Ligase PafA. *Journal of the American Chemical Society* **135**, 6794-6797,(2013).

3 Castano-Cerezo, S. *et al.* Protein acetylation affects acetate metabolism, motility and acid stress response in Escherichia coli. *Molecular Systems Biology* **10**,(2014).

4 Weinert, B. T. *et al.* Acetyl-Phosphate Is a Critical Determinant of Lysine Acetylation in E. coli. *Molecular Cell* **51**, 265-272,(2013).
